# Speeding up anterior-posterior patterning of insects by differential initialization of the gap gene cascade

**DOI:** 10.1101/485151

**Authors:** Heike Rudolf, Christine Zellner, Ezzat El-Sherif

## Abstract

Recently, it was shown that anterior-posterior patterning genes in the red flour beetle *Tribolium castaneum* are expressed sequentially in waves. However, in the fruit fly *Drosophila melanogaster*, an insect with a derived mode of embryogenesis compared to *Tribolium*, anterior-posterior patterning genes quickly and simultaneously arise as mature gene expression domains that, afterwards, undergo slight posterior-to-anterior shifts. This raises the question of how a fast and simultaneous mode of patterning, like that of *Drosophila*, could have evolved from a rather slow sequential mode of patterning, like that of *Tribolium*. In this paper, we elucidate a mechanism for this evolutionary transition based on a switch from a uniform to a gradient-mediated initialization of the gap gene cascade by maternal Hb. The model is supported by computational analyses and experiments.

## Introduction

A common problem in development is how to partition a tissue into serial structures and/or regions of different fates. Interestingly, the generation of serial structures, in many cases, was found to rely on oscillatory gene activity mediated by molecular clocks (e.g. vertebrate somitogenesis (1-3) and lateral roots specification in plants (4, 5)). Similarly, the division of a tissue into different (aperiodic) fates was found to rely on aperiodic sequential activation of genes mediated by genetic cascades (e.g. *Drosophila* neurogenesis (6, 7), vertebrate neural tube patterning (8), and anterior-posterior (AP) fate specification in vertebrates (9) and short-germ insects (10, 11)).

Spatial patterning is usually mediated by a morphogen gradient, secreted from a signaling center or an organizer. In cases where temporal rhythms are involved in patterning, genes are usually expressed in spatial waves that emanate and propagate away from the organizer (Figure 1). These waves eventually stabilize into gene expression domains. For example, during vertebrate somitogenesis, Wnt and FGF form posterior-to-anterior gradients in the presomitic mesoderm. Oscillatory expression waves of the transcription factor family *hes*/*hairy* and components of several signaling pathways were observed to emanate from the posterior end of the presomitic mesoderm and propagate away to form striped expression more anteriorly (Figure 1D, left), delimiting vertebrate somites (1, 12-15). In parallel, sequential waves of Hox gene expressions propagate from posterior to anterior to partition the vertebrate AP axis into different fates (Figure 1D, right) (9, 16, 17). During neural tube patterning in vertebrates, Shh forms a ventral to dorsal gradient. Neural fates emanate sequentially from ventral and spread towards more dorsal regions of the neural tube in the high-to-low directions of the Shh gradient (Figure 1B) (8, 18–20). Similar observations were made in other developmental systems (e.g. lateral root formation in plants (4, 5) and limb bud patterning in vertebrates (21-23); Figures 1 A and C, respectively). Hence, patterning by means of gene expression waves seems to be a prevalent mechanism in development.

**Figure 1.**
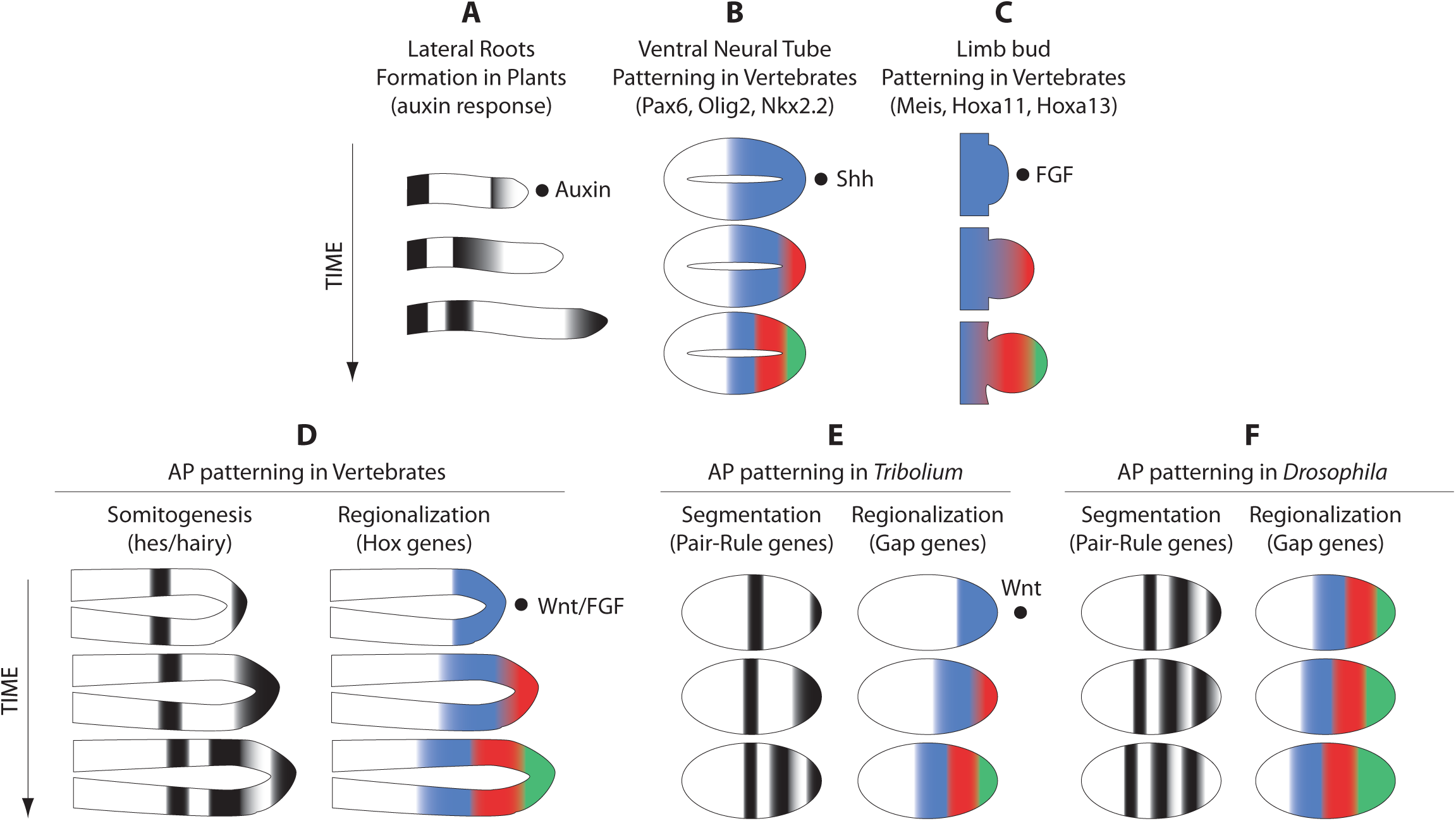
Gene expression waves in development. Many embryonic structures are patterned via periodic and/or aperiodic waves of gene expressions that emanate from an organizer. **(A)** Periodic waves of _ emanate from an Auxin organizer to demarcate lateral roots. **(B)** Aperiodic waves of gene expression emanate from a ventral organizer secreting Shh ligands to pattern the ventral neural tube in insects. Ventral to the right. **(C)** Aperiodic waves of gene expressions emanate from a distal organizer secreting FGF ligands to pattern the limb bud. Distal to the right. **(D)** Periodic waves of *hes*/*hairy* expression and aperiodic waves of Hox genes expressions emanate from a posterior organizer secreting Wnt/FGF ligands to segment and regionalize the anterior-posterior axis of vertebrates, respectively. **(E)** Periodic waves of pair-rule genes expressions and aperiodic waves of gap genes expressions emanate from a posterior organizer secreting Wnt ligands to segment and regionalize the anterior-posterior axis of short-germ insects, respectively. **(F)** Pair-rule and gap gene expressions arise simultaneously in the *Drosophila* embryo, but then exhibit vestiges of wave dynamics in the form of posterior-to-anterior shifts. Posterior to the right in (D, E, and F).

An interesting case, however, is the AP patterning of the fruit fly *Drosophila melanogaster*. The AP axis of *Drosophila* (and most insects) is divided into different fates by the spatially aperiodic expressions of a group of genes called ‘gap genes’, and into repeating units (segments) by the spatially periodic expressions of a group of genes called ‘pair-rule genes’ (24, 25). The expressions of both gap and pair-rule genes in *Drosophila* seem initially to arise simultaneously and *de novo* as mature gene expression domains, with no apparent sequential/oscillatory temporal dynamics (Figure 1F) (25). However, later on, both gap and pair-rule gene expression domains undergo posterior-to-anterior shifts, reminiscent of gene expression wave dynamics (Figure 1F) (25-29). Interestingly, in the beetle *Tribolium castaneum* (which is thought to adopt an ancestral mode of AP patterning), both gap and pair-rule genes are expressed in outright sequential/oscillatory waves (Figure 1E) (10, 30, 31). Hence, the simultaneous mode of gap and pair-rule gene regulation in *Drosophila* seems to have evolved from a more ancestral mode of sequential/oscillatory gene regulation, bearing vestiges of the ancestral mode in the form of posterior-to-anterior shifts of gene expression domains.

Another important difference between the ancestral mode of AP patterning in insects (arguably exemplified by *Tribolium*) and that of *Drosophila* is the morphology of the embryo at the time of AP fate determination. The AP axis of most insects is partitioned into different fates in two different phases, each with a different morphology (Figure 2A). First, anterior fates arise in a ‘blastoderm’, a structure with a fixed AP length. Then, more posterior fates form in a ‘germband’, whose AP axis grows by convergent extension and/or cell division. Insects differ in the number of fates that form in the blastoderm vs germband (32). In short-germ insects (e.g. the grasshopper *Schistocerca Americana* (33)), most fates form in a germband; while in long-germ insects (e.g. the wasp *Nasonia vitripennis* (34), the fruit fly *Drosophila melanogaster*, and the bean beetle *Callosobruchus maculatus* (35)), most fates form in a blastoderm. Intermediate-germ insects (e.g. the flour beetle *Tribolium castaneum*, the milkweed bug *Oncopeltus fasciatus* (36, 37), and the beetle *Dermestes maculatus* (38)) lie somewhere in between, where anterior fates form in the blastoderm and posterior fates form in the germband stage (Figure 2A). Short-germ embryogenesis is thought to be the ancestral mode of insect development, and an evolutionary trend of short-germ to long-germ evolution is observed to occur independently several time throughout evolution (with some reports of the opposite path (39)).

**Figure 2.**
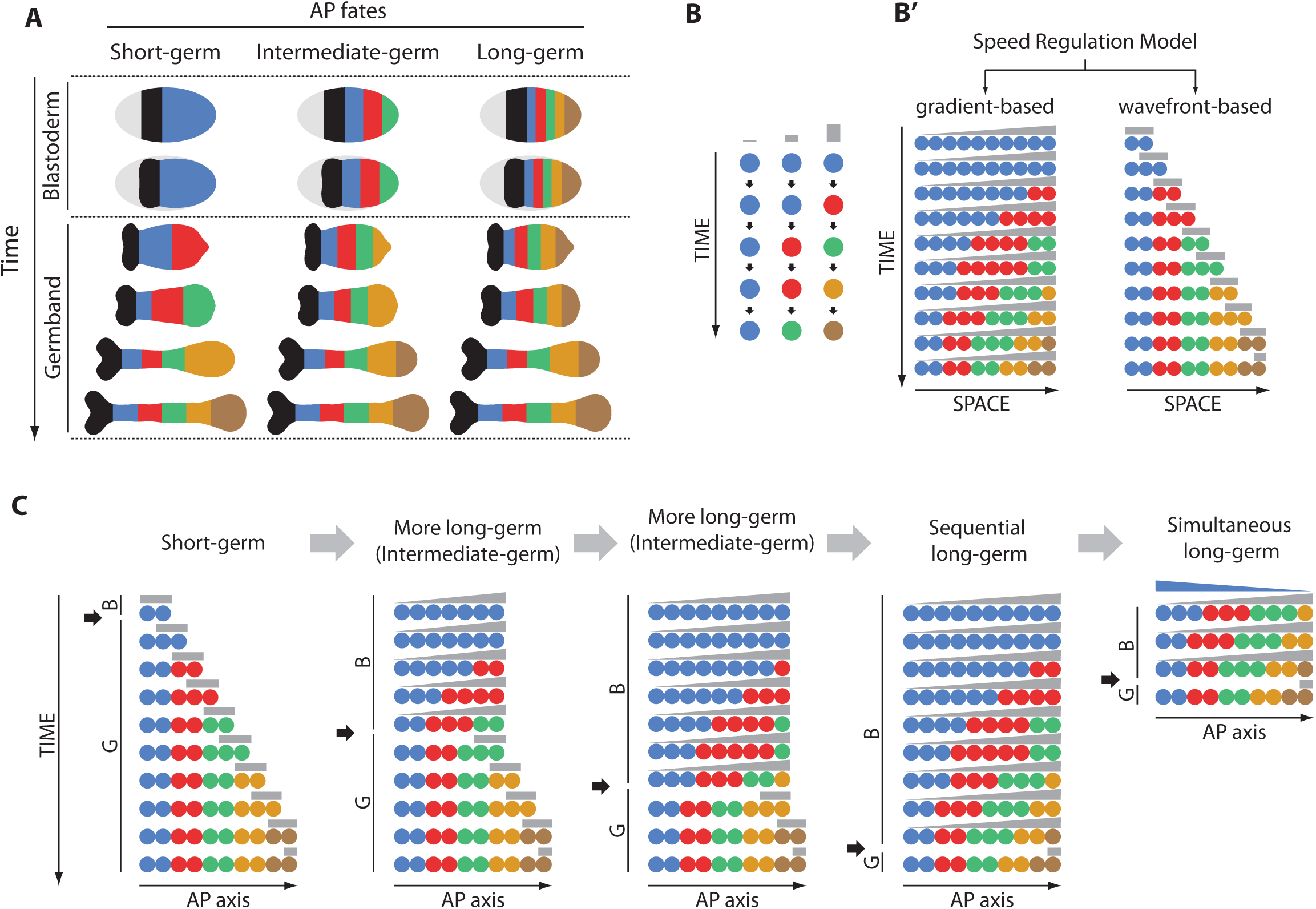
A Speed Regulation model for short- to long-germ evolution. **(A)** In short-germ insects, most of AP fates (shown in blue, red, green, gold, and brown; black marks the head; grey marks the extraembryonic tissue) are specified in the germband stage. In intermediate-germ insect, anterior fates are specified in the blastoderm while posterior fates are specified during the germband stage. In long-germ insects, most fates are specified in the blastoderm. **(B, B’)** In the Speed Regulation model, a molecular factor (speed regulator; shown in grey) regulate the speed of the sequential activation of fates (shown in different colors) (B). The Speed Regulation model can operate in a gradient-based mode to pattern non-elongating tissues (B’, left), or in a wavefront-based mode to pattern elongating tissues (B’, right). **(C)** The Speed Regulation model can explain the evolution of sequential short-germ embryogenesis to simultaneous long-germ embryogenesis. Can it also explain simultaneous long-germ embryogenesis?

Insects then differ in their modes of AP patterning in two different (but possibly related) aspects: (1) whether they undergo short-, intermediate-, or long-germ embryogenesis, and (2) whether AP fates are determined sequentially or simultaneously. Are these two aspects correlated? Sequential patterning is naturally more suitable for an elongating tissue like the germband (which indeed is the case in the germbands of most insects). Simultaneous patterning, on the other hand, is expected to be limited to a non-elongating tissue like the blastoderm (which indeed is the case in the blastoderm of the lon-germ insect *Drosophila*). It is conceivable, nonetheless, that a blastoderm would be patterned sequentially. Indeed, the blastoderm of the intermediate-germ insect *Tribolium* is patterned by a sequential/oscillatory mechanism (10, 31). Examining the mode of AP patterning in more insects is needed to give a definite answer to this question. However, based on these (rather few) study cases, one is tempted to extrapolate that short-germ and intermediate-germ insects tend to adopt a sequential patterning mechanism in both their blastoderm and germband stages, whereas simultaneous patterning is more commonly found in the blastoderms of long-germ insects.

Environmental pressure for fast patterning is potentially the major cause for the evolution of sequential to simultaneous patterning (for example, AP patterning is complete after 3 hours post fertilization in *Drosophila* and after 48 hours in *Tribolium*). However, the mechanism behind this sequential-to- simultaneous evolution is not clear. In this paper, we present a simple model for sequential-to- simultaneous evolution of gap gene regulation in insects. In this model, simply expressing the first gene in the gap gene cascade (namely, *hunchback*, *hb*) in an anterior-to-posterior gradient speeds up the formation of gap gene domains. Expressing *hb* in yet steeper gradient further speeds up patterning and ultimately leads to almost simultaneous emergence of gap gene domains. The model recapitulates key experimental observations in *Drosophila*. Furthermore, the effect of experimentally manipulating the Hb gradient in *Tribolium* is consistent with our model.

### A model for the evolution of sequential short-germ to simultaneous long-germ embryogenesis in insects

A model for <u>sequential</u> *short-germ* to <u>sequential</u> *long-germ* evolution has been recently devised based on a Speed Regulation model of embryonic patterning (Figure 2B, B’, C) (10, 40). In this model, each cell in an embryonic structure has the capacity to transit through successive states (shown in different colors in Figure 2B). The speed of state transitions is regulated by a molecular factor (shown in grey in Figure 2B, and henceforth called a ‘speed regulator’). If a group of cells is subject to a gradient of the speed regulator (Figure 2B’, left), all cells go through successive states, but with slower and slower speed as we go from higher to lower values of the speed gradient. This gives the appearance that the states propagate as waves in the high-to-low direction of the gradient (Figure 2B’, left). These waves do not require diffusion or cell-cell communication and, hence, are called “kinematic” or “pseudo-waves” (1, 13, 30, 31, 41–44). The “Speed Regulation” model (Figure 2B, B’) can pattern tissues with no or limited axial elongation using the gradient-based mode of the model (Figure B’, left). It can also pattern elongating tissues if the gradient is retracting as a wavefront (the wavefront-based mode of the Speed Regulation model, Figure 2B’, right) (10). Gene expression waves can also be generated in the wavefront-based mode if the retracting gradient of the speed regulator is tapered, as indeed observed during vertebrate somitogenesis (13).

Since the Speed Regulation model can pattern both non-elongating tissues (like the blastoderm) and elongating tissues (like the germband), it is easy to imagine an evolutionary scenario where germband fates shift into the blastoderm by simply delaying blastoderm-to-germband transition (Figure 2C), or, alternatively, speeding up fate transitions so that most fates form in the blastoderm. In this case, AP fates (specified by gap genes) are expected to be expressed sequentially in the blastoderm of intermediate-germ insects (which indeed is the case in *Tribolium*) (10) and long-germ insects. However, in case of the long-germ insect *Drosophila*, gap genes seem to be initially activated simultaneously, and then undergo slight posterior-to-anterior shift (26, 27). This behavior cannot be explained by our current formulation of the Speed Regulation model. In addition, the model assumes that AP patterning is mediated primarily by a posterior-to-anterior morphogen gradient, which is the case in short/intermediate-germ insects like *Tribolium*, but definitely not the case in *Drosophila*. Indeed, gap gene regulation in *Tribolium* seems to be primarily mediated by a posterior-to-anterior gradient of Wnt/Caudal (10). Depleting *caudal* (*cad*) transcripts by RNAi completely abolishes trunk gap gene expressions. Furthermore, the gap gene cascade can be re-induced in the *cad*-expressing posterior end of the embryo in the germband stage, where the influence of anterior-to-posterior gradients is unlikely (45). On the other hand, the gap gene system in *Drosophila* is heavily dependent on anterior-to-posterior morphogen gradients (namely, Bicoid and maternal Hb) and is less dependent on the posterior-to-anterior gradient of Cad (24, 25, 46, 47). We then wondered if the adoption of a patterning mechanism based on Bcd and maternal Hb gradients could have mediated the evolution of a sequential regulation of gap genes to a more simultaneous mechanism (Figrue 2C, last column). Since *bcd* is lacking in non-Dipterans, and since *bcd* (48-50) and maternal Hb seem to act redundantly in regulating gap genes in *Drosophila* (51), we narrowed down our search for a potential evolutionary bridge between sequential and simultaneous patterning to the maternal Hb gradient.

We note here that *hb* is special in that it is the first gene in the trunk gap gene cascade in insects (blue gene in Figure 2) (24). In the Speed Regulation model, cells along the spatial axis are all initialized with the same concentration of the blue gene (Figure 2B, B’, C). What happens then if we initialize cells with a gradient of the blue gene expression (analogous to a maternal Hb gradient)? Would this lead to a *Drosophila*-like simultaneous mode of patterning (last column in Figure 2 C)?

Since gene expressions are modeled as either on or off in the Speed Regulation model, the model cannot predict the outcome if initialized with graded values of gene expressions. Hence, we use a gene network realization of the Speed Regulation model recently suggested in refs (11, 12), namely: the gradual module switching model (10, 40) (Supplementary Text S1). We first used the module switching scheme to model the AP regionalization in a short-germ insect (Figure 3A; Movie S1A), where very few AP fate-determining genes (the blue gene and partially the red gene in Figure 3A) are expressed during the blastoderm stage. During the germband stage, the rest of genes (red, green, gold, and brown in Figure 3, first column) start to emanate sequentially from posterior (Figure 3A; Movie S1A).

**Figure 3.**
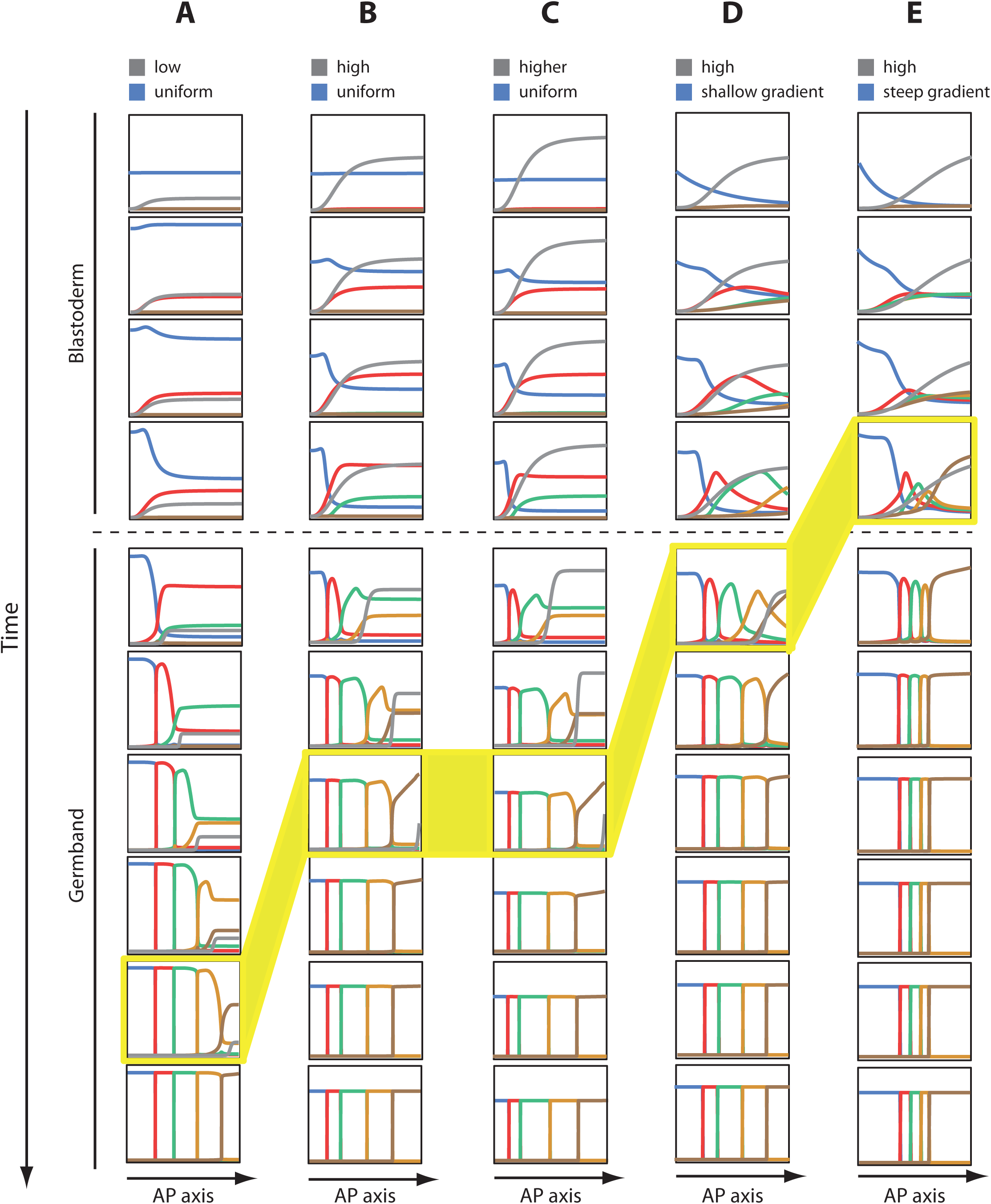
Effect of changing posterior morphogen concentration and first gene initialization on patterning speed in a molecular realization of the Speed Regulation model. **(A)** In a molecular realization of the Speed Regulation principle (in particular, the Gradual Module Switching model) to model short-germ embryogenesis, few anterior fate-specifying gene expression bands (here blue and partially red) form in the blastoderm, while the rest form in the germband (partially red, green, gold, and brown). The speed regulator (shown in grey) is expressed in a static posterior-to-anterior gradient in the blastoderm, while it is expressed in a posteriorly retracting wavefront in the germband. The timepoint at which all gene expression bands arise is highlighted in yellow. **(B)** Increasing the concentration of the speed regulator (grey) speed up the patterning process. Note the timepoint at which all expression bands arise (highlighted in yellow); compare with (A). **(C)** Further increasing the concentration of the speed regulator does not result in a further increase in patterning speed. Note that in A, B, and C, the first gene in the cascade (the blue gene) is expressed uniformly along the AP axis. **(D)** Expressing the blue gene as a shallow anterior-to-posterior gradient speeds up the patterning process further; however, gene expression patterns still arise sequentially. **(E)** Expressing the blue gene as a steep anterior-to-posterior gradient speeds up the patterning process even further. Gene expression domains now arise simultaneously. Posterior to the right. Timepoints at which all gene expression bands form are highlighted in yellow.

We then sought to evolve our short-germ embryogenesis model (Figure 3A) into a more intermediate/long-germ one. One strategy is to delay the blastoderm-to-germband transition (Figure 2C). Another strategy is to speed up the genetic cascade so that more fates are determined during the blastoderm and before the blastoderm-to-germband transition. Here we will follow the latter strategy since AP patterning in long-germ insects like *Drosophila* tend to be faster than in short- and intermediate-germ insects. Since the posterior morphogen is acting as a speed regulator in our model, a natural strategy to speed up AP patterning would be to increase the concentration of the speed regulator. Indeed, increasing the level of the posterior morphogen sped up AP patterning (Figure 3B; Movie S1B), and now more AP fates form during the blastoderm stage. This strategy has a limit, however, since further increasing the level of the posterior morphogen does not result in further increase in the speed of AP patterning (Figure 3C, Movie S1C), possibly due to a saturation effect in the module switching model.

So far in our simulations we initialized the AP axis with a uniform expression of the first gene in the cascade (blue gene in Figure 3 and Movie S1). We then expressed the blue gene in an anterior-to-posterior gradient (in a similar fashion to the anterior-to-posterior Hb gradient in *Drosophila*). Surprisingly, this further sped up patterning (Figure 3D, Movie S1D), resulting in more fates forming during the blastoderm stage, and hence mimicking a further evolution into long-germ embryogenesis. However, the generation of gene expression domains, albeit faster, was still sequential. We then sought to apply a steeper gradient of the first gene (Figure 3E, Movie S1E). This, surprisingly, led to speedy patterning where gene expression domains arouse simultaneously during the blastoderm stage, in accord with the fast and simultaneous patterning of gap genes in *Drosophila*.

### How does differential initialization of genetic cascades result in fast simultaneous patterning?

Thus, simply expressing the first gene in the cascade in an anterior-to-posterior gradient resulted in a fast and simultaneous patterning in the context of Speed Regulation model (or rather its gene network realization: the module switching model). However, it is not clear what factors contributed to this effect. To investigate this, we traced the temporal evolution of the model outputs at different positions along the AP axis for two different cases: (*i*) when the AP axis is initialized with a uniform expression of the first gene (Figure 4 A), and (*ii*) when the AP axis is initialized with an anterior-to-posterior steep gradient of the expression of the first gene (Figure 4 B). In both cases, genes are activated sequentially. However, in the latter case, genes are activated only weakly and transiently before a specific gene in the cascade is fully activated. Which gene is fully activated (after previous genes are transiently and weakly activated) in this case depends on the starting concentration of the first gene, and hence, on spatial position. This behavior, hence, results in fast activation of genes and gives the impression of simultaneous patterning. Interestingly, uniformly expressing the speed regulator (grey in Figure 4) across space but expressing the first gene in the cascade in an anterior-to-posterior gradient resulted in a fast and simultaneous patterning as well (Figure 4 C, D; Movie S2). Hence, expressing the first gene in the cascade as a gradient is enough to establish the full expression pattern, without the need for the speed regulator to be expressed in a gradient.

**Figure 4.**
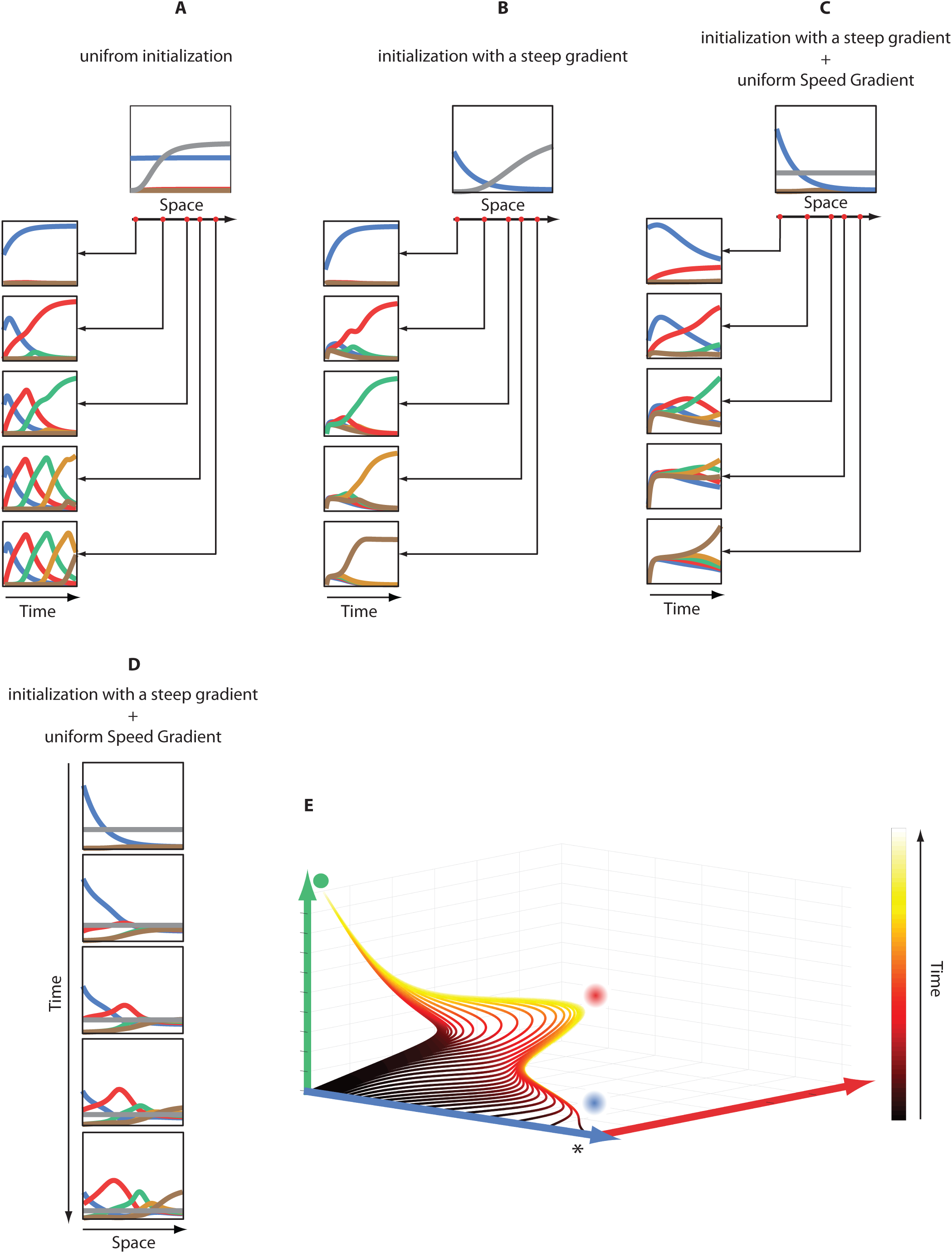
How differential initialization of genetic cascades mediates simultaneous patterning. **(A)** Time courses of the Gradual Module Switching model at different positions along the AP axis if the speed gradient (grey) is expressed in a posterior-to-anterior gradient, and the first gene in the cascade (blue) is expressed uniformly. **(B)** Time courses of the Gradual Module Switching model at different positions along the AP axis if the speed gradient (grey) is expressed in a posterior-to-anterior gradient, and the first gene in the cascade (blue) is expressed in an anterior-to-posterior gradient. **(C)** Time courses of the Gradual Module Switching model at different positions along the AP axis if the speed gradient (grey) is expressed uniformly, and the first gene in the cascade (blue) is expressed in an anterior-to-posterior gradient. **(D)** The spatiotemporal dynamics of the Gradual Module Switching model outputs if the speed gradient (grey) has a uniform (but decaying) expression along space, and the first gene in the cascade (blue) is expressed in an anterior-to-posterior gradient. **(E)** State space representation of a 3-genes genetic cascade with different initial concentrations of the first gene in the cascade (blue gene). Solid green circle: fixed point attractor of the whole system, characterized by high concentration of the green gene and low concentrations of the blue and red genes. Fuzzy blue circle: a ghost attractor characterized by high concentration of blue gene expression and low concentrations of the red and green gene expressions. Fuzzy red circle: a ghost attractor characterized by high concentration of red gene expression and low concentrations of the blue and green gene expressions

To gain more insight into how differentially initializing a genetic cascade along space leads to seemingly simultaneous patterning, we generated a phase space diagram (Figure 4E) for the outputs of a 3-genes genetic cascade initialized with different initial concentrations of the first genes (with zero initial concentration of the second and third genes). The phase space in Figure 4E contains one fixed point attractor (green fixed point). The ‘green fixed point’ is characterized by a high concentration of the green gene expression (last gene in the 3-genes cascade) and low concentration of the blue and red gene expressions (Figure 4E). Additionally, the phase space contains two of what are called ‘ghost attractors’ (52, 53) (fuzzy blue and fuzzy red circles in Figure 4E), which are not real fixed points at which the system can stay at steady state, but are points of attraction, nonetheless. The blue ghost attractor is characterized by a high level of the blue gene expression but low levels of red and green genes expressions. The red ghost attractor is characterized by a high level of the red gene expression but low levels of blue and green genes expressions. When the genetic cascade is initialized with a high concentration of the first (blue) gene expression (asterisk in Figure 4E), the output follows a trajectory expected from a typical genetic cascade: the system first approaches the blue ghost attractor, and then approaches the red ghost attractor, and finally reaches the green fixed point (the final gene in the cascade). However, upon progressively lowering the initial concentration of the blue gene expression, the trajectory of the system progressively skips the blue ghost attractor and heads directly to the red ghost attractor, before finally reaches the green fixed point. Upon lowering the initial concentration of the blue gene expression further, the trajectory even skips both blue and red ghost attractors and heads (almost) directly to the green fixed point. This ‘ghost attractor skipping’ behavior then is the cause of the fast and seemingly simultaneous patterning.

### Experimental support in *Drosophila*

Thus, a patterning mechanism based on the speed regulation of a genetic cascade by a posterior-to-anterior morphogen (like that hypothesized for ancestral insects) can smoothly evolve into a simultaneous patterning mechanism based (possibly solely) on an anterior-to-posterior gradient of the first gene in the cascade. In refs (10, 45), we provided evidence that patterning the AP axis of *Tribolium* is based on the former mechanism. Here we argue that patterning the AP axis of *Drosophila* is (partially) based on the latter mechanism.

The partitioning of the AP axis of *Drosophila* (excluding the head region) into different fates is mediated by the expressions of four gap genes: *hb*, *Krüppel* (*Kr*), *knirps* (*kni*), and *giant* (*gt*) (Figure 5A; an additional posterior domain of *hb* and a head expression of *gt* are not shown and excluded from our analysis as their expressions are mediated by the terminal system in *Drosophila*) (24, 25). Three maternally provided morphogen gradients were shown to be involved in regulating these four gap genes: (*i*) the anterior-to-posterior gradient of Bcd (54, 55), (*ii*) the anterior-to-posterior gradient of maternal Hb (51, 56), and (*iii*) the posterior-to-anterior gradient of Cad (57). During oogenesis, *nanos* (*nos*) (58) and *bcd* mRNAs (54) are localized to the posterior and anterior poles of the *Drosophila* embryo, respectively, while *hb*, *pumilio* (*pum*) (59), and *cad* mRNAs are ubiquitously distributed (57, 60). During embryogenesis, *bcd* is translated into an anterior-to-posterior Bcd gradient. Bcd gradient then translationally represses the ubiquitously expressed *cad* mRNAs forming a posterior-to-anterior Cad gradient (60) (in addition to seemingly redundant zygotic activation of *cad* (61)). The posteriorly localized *nos* is translated into a posterior-to-anterior gradient of *Nos* proteins. Nos together with Pum and other factors form a repression complex that binds Nanos response element (NRE) in the 3′ UTR of the *hb* mRNAs and mediate their deadenylation (62). This results in an anterior-to-posterior gradient of Hb proteins. In addition, *hb* is zygotically regulated as part of the gap gene network. Thus, Hb protein distribution progressively turns from an early maternal gradient to a zygotically regulated anterior domain.

**Figure 5.**
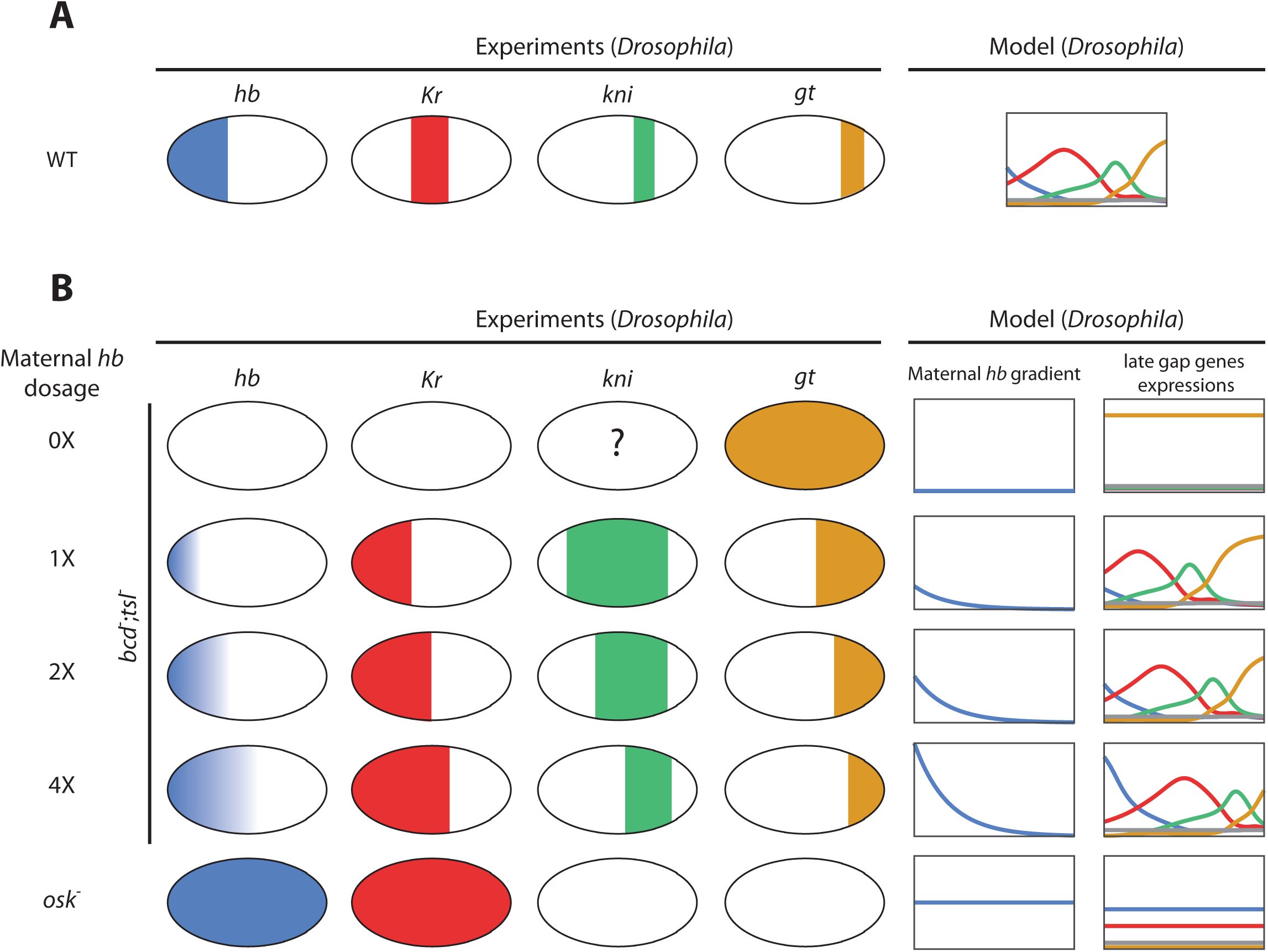
Model versus Experiment: Effect of varying the dosage of maternal Hb in *Drosophila*. **(A)** *Drosophila* gap gene domains in WT: experiments (left) versus our model (right). **(B)** Increasing the dosage of maternal Hb in *bcd*;*tsl* background (in addition to an *osk* mutant in which maternal Hb is distributed uniformly in the embryo; last row) leads to progressive posterior shifts of gap gene expression domains in both experiment (left) and a model of gap gene regulation in *Drosophila* (right).

The anterior-to-posterior gradients of Bcd and maternal Hb were shown to provide positional information for the gap gene expression domains in *Drosophila* (54, 56, 63). However, complete absence of either gradients alone results only in an anterior shift of the gap gene domains (51). Embryos lacking both Bcd and maternal Hb, however, show severe disruption of gap gene regulation (51), indicating that Bcd and maternal Hb are acting redundantly (partially because Bcd activates *hb*). In the absence of Cad (from both maternal and zygotic contributions), only weak effects were observed on gap gene expressions (46). In the absence of both Bcd and Cad, however, gap gene expressions are severely disrupted, indicating a redundancy between these two gradient systems (57).

Thus, three morphogen gradients (Bcd, maternal Hb, and Cad) are acting (redundantly) to regulate gap genes in *Drosophila*. Since *bcd* is an evolutionary novelty specific to Diptera, the maternal Hb and Cad gradients are reasonable candidates for an ancestral AP morphogen system. As mentioned earlier, maternal Hb gradient seems sufficient to provide positional information for gap gene expressions in the *Drosophila* embryo. Embryos mutants for both *bcd* and *torso-like* (*tsl*) lack a Bcd gradient and gradients generated by the terminal system. The lack of Bcd results in a ubiquitous distribution of maternal Cad proteins (with a possible weak zygotic contribution at the posterior). Thus, in *bcd*;*tsl* mutant embryos, maternal Hb is (almost) the only maternal gradient left in the *Drosophila* embryo (56). In these embryos, the correct order of gap genes is generated: *hb*, *Kr*, *kni*, and *gt* (listed from anterior to posterior). In a classical study ((56); which results are reproduced in Figure 5B, experiments), the dosage of maternal *hb* was manipulated in *bcd*;*tsl* embryos. In addition, a mutation in the gene *oskar* (*osk*) to eliminate *nos* activity was used to generate embryos with uniform distribution of Hb proteins throughout the embryo (56, 64). Examining gap gene expressions in these embryos confirmed that the maternal Hb gradient specifies the positions of gap gene domains along the AP axis in a dose dependent manner (56). In particular, a progressive increase in maternal Hb dosage leads to a progressive shift in the gap gene domains towards the posterior pole of the embryo. This effect can be recapitulated using our model for simultaneous patterning. A computational model in which *Drosophila* gap genes (*hb*, *Kr*, *kni*, and *gt*) are wired into a genetic cascade and initialized by a maternal gradient of Hb (first gene in the cascade) generated the correct order of gap genes expression domains along the AP axis (compare model to experiments in Figure 5A). Manipulating the dosage of Hb in our model led to similar shifts in the positions of gap gene expression domains as observed experimentally (Figure 5B: compare model to experiments; Movie S3).

We then sought to examine if our model can predict the outcome of further genetic perturbations in *Drosophila*. In Figure 6 (experiments), we summarized published expression patterns of different gap genes in WT and various gap gene mutants in *Drosophila*. Examining these results, we noticed a general pattern: the absence of the expression of one gene led to the downregulation of the gene immediately posterior to it. The bands anterior and posterior to the missing gene expression domains extend (towards posterior and anterior, respectively) to take up the space normally occupied by the missing bands in WT. In *Kr*-embryos, *kni* expression is missing (65, 66), while anterior *hb* expression extends posteriorly (67-69), and posterior *gt* expression extends anteriorly (70, 71) (compare Figure 6B to 6A, experiments). In *kni*-embryos, *gt* expression is weak (70, 72), while *Kr* expression extends posteriorly (67) (compare Figure 6C to 6A, experiments). In *gt*-embryos, *kni* expression extends posteriorly (72) (compare Figure 6D to 6A, experiments). The downregulation of one gene upon quenching the expression of the gene anterior to it is intriguingly analogous to the behavior of a genetic cascade: the activation (or de-repression) of a gene in the cascade is dependent on the activity of the gene preceding it in the cascade. Indeed, our model for gap gene regulation in *Drosophila* (which is based on a genetic cascade of gap genes) recapitulated all experimental gap gene perturbations (Figure 6; compare model to experiments; Movie S4).

**Figure 6.**
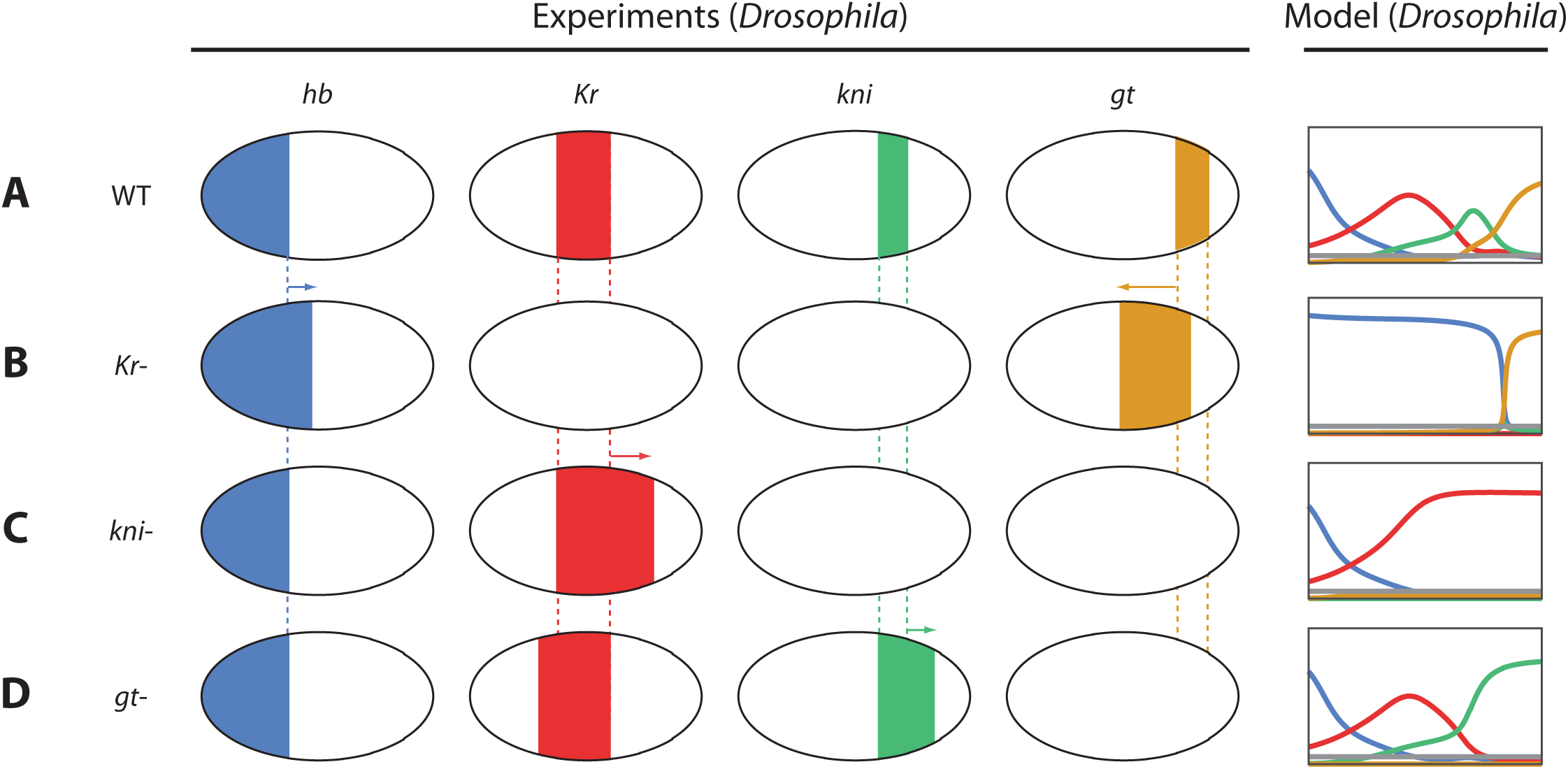
Model versus Experiment: gap mutants in *Drosophila*. Expression patterns of *Drosophila* gap genes in WT and gap gene mutants in both experiment (left) and of gap gene regulation in *Drosophila* (right).

### Experimental support in *Tribolium*

So far, we have argued that using maternal Hb gradient to regulate the gap gene cascade is an important evolutionary link in the evolution of sequential to simultaneous patterning of the AP axis in insects. Does a maternal Hb gradient, then, already exist in sequentially patterning insects? Indeed, Hb proteins were found to have an anterior-to-posterior graded distribution along the AP axis of early *Tribolium* blastoderm, albeit more extended towards posterior compared to the maternal Hb gradient in *Drosophila* (Figure 7 A, A’). The Hb gradient in *Tribolium* was shown to be mediated, like *Drosophila*, by *nos* and *pum*, as Hb proteins were found to be uniformly distributed in the early *Tribolium* blastoderm in *nos*;*pum* double RNAi (Figure 7A) (73). But what could be the function of a *nos*/*pum*-mediated maternal Hb gradient in the early embryo of a sequentially segmenting insect like *Tribolium*? As shown in Figure 3, initially expressing Hb even as a shallow gradient could speed up patterning (Figure 3D), albeit not achieving a simultaneous mode of patterning (which could be achieved using a steep Hb gradient, Figure 3E, as indeed the case in *Drosophila*; compare Figure 7 A’ to 7 A).

**Figure 7.**
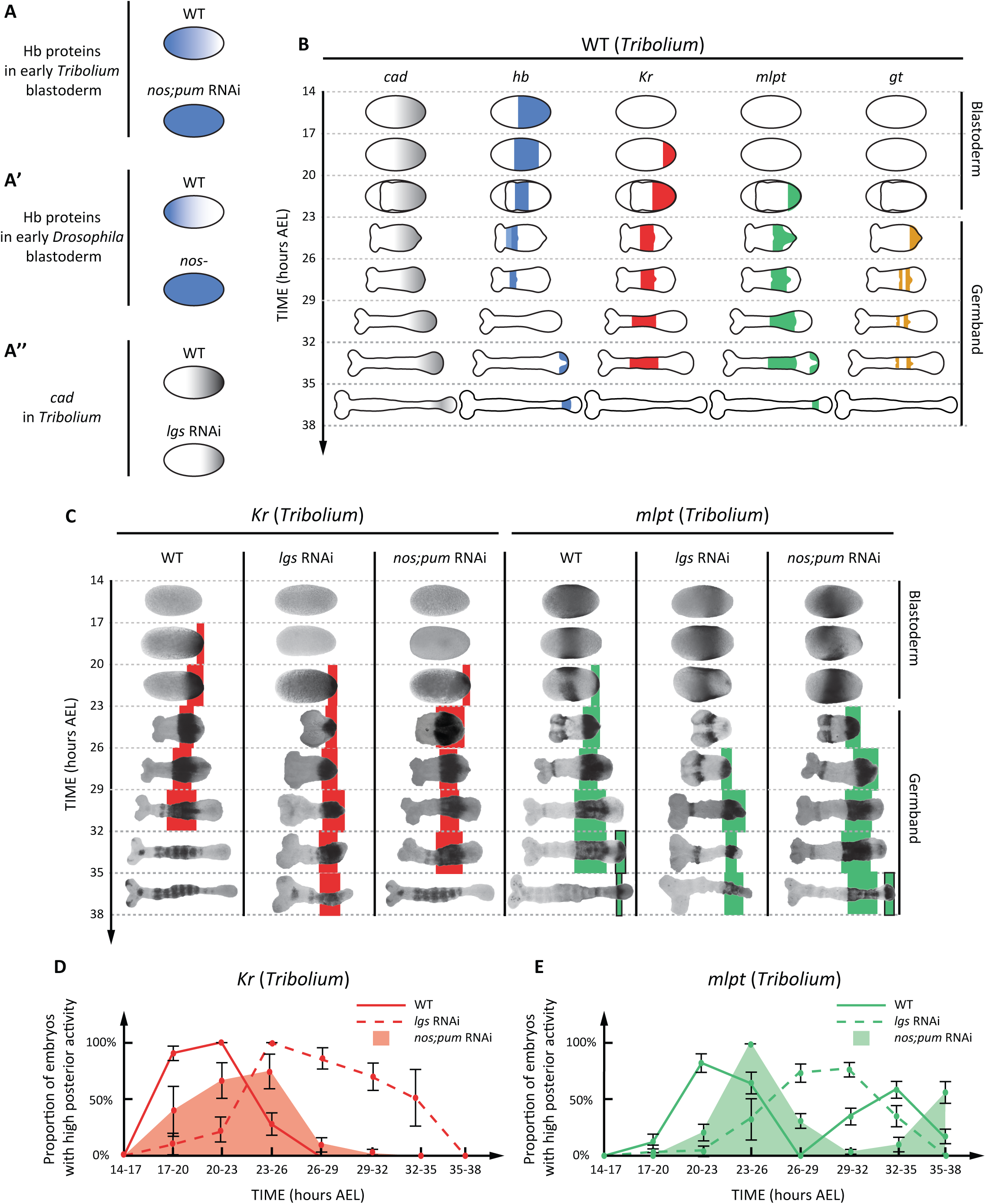
The role of maternal Hb gradient in the AP patterning of *Tribolium*. **(A)** Hb proteins are distributed in an anterior-to-posterior gradient in early WT *Tribolium* blastoderm. Hb gradient becomes uniform in *nos*;*pum* RNAi embryos. **(A’)** The maternal Hb gradient in *Drosophila* is shifted more to the anterior compared to *Tribolium*. **(A’’)** *cad* gradient is reduced and shifted to the posterior in *lgs* RNAi *Tribolium* blastoderm compared to WT. **(B)** The spatiotemporal dynamics of *cad*, *hb*, *Kr*, *mlpt*, and *gt* in *Tribolium*. **(C)** The spatiotemporal dynamics of *Tribolium Kr* and *mlpt* in WT, *lgs* RNAi, and *nos*;*pum* RNAi embryos. **(D)** Quantification of the temporal dynamics of *Kr* at the posterior end of WT, *lgs* RNAi, and *nos*;*pum* RNAi *Tribolium* embryos. **(E)** Quantification of the temporal dynamics of *mlpt* at the posterior end of WT, *lgs* RNAi, and *nos*;*pum* RNAi *Tribolium* embryos.

Hence, we wondered if the maternal Hb gradient in *Tribolium* helps speeding up AP patterning in the *Tribolium* blastoderm. In *Tribolium*, the expression domains of four gap genes (*hb*, *Kr*, *milles-pattes* (*mlpt*), and *gt*) (24, 74-79) emanate sequentially in waves from a posterior Active Zone (AZ), where *cad* is expressed in a posterior-to-anterior gradient (see the data summary cartoon in Figure 7B) (10). Gap gene expression waves propagate anteriorly and freeze into stable domains upon exiting the AZ. It was previously suggested that gap gene expression waves are mediated by a Speed Regulation mechanism where *cad* is acting as a speed regulator. The mechanism is applicable to both blastoderm and germband stages as *cad* is expressed in a posterior-to-anterior gradient in both stages. In contrast, Hb is expressed in an anterior-to-posterior gradient only in early blastoderm stage. Hence, any potential role of maternal Hb in speeding up patterning should be a transient one during the blastoderm stage, specifically involved in regulating the blastodermal expressions of *hb*, *Kr*, and *mlpt* (Figure 7B; blastoderm stage: 14-23 hours After Egg Lay (AEL)).

Hence, according to our models, there exists two factors that influence the speed of gap gene expression sequence: concentration of *cad* and the maternal Hb gradient. Either reducing *cad* or expressing maternal Hb uniformly (rather than in a gradient) is expected to reduce the speed of the gap gene expression sequence. However, the effect of maternal Hb on patterning speed is expected to be transient during the blastoderm stage, while the effect of *cad* is expected to be persistent in both blastoderm and germband stages.

To test our predictions for these two cases, we examined the spatiotemporal dynamics of two of the gap genes (*Kr* and *mlpt*) in *Tribolium* for: WT embryos, embryos where *cad* concentration is reduced (by knocking down the Wnt positive regulator *legless* (*lgs*) using RNAi (10, 30, 80–82)), and embryos where maternal Hb gradient is made uniform (*nos*;*pum* RNAi knockdown embryos) (Figure 7C). For more quantitative analysis, we examined the statistics of the temporal dynamics of *Kr* and *mlpt* expressions in the posterior end of the *Tribolium* embryo (where the sequential activation of gap genes is of the highest speed) for WT, *lgs* RNAi, and *nos*;*pum* RNAi embryos (Figure 7D, E; Methods). In ref (10), we already carried out a similar analysis for *lgs* RNAi embryos. However, the analysis was restricted to the initial phase of patterning (mainly blastoderm stage), and we are here extending the analysis to the germband stage as well.

In WT embryos, *Kr* expression emanates from the posterior end of the blastoderm roughly at 17-20 hours AEL and clears from posterior at 23-26 hours AEL during the germband stage (Figure 7C, D). In *lgs* RNAi embryos, the initiation of *Kr* expression is delayed, starting around 20-23 hours AEL (Figure 7C, D). *Kr* expression then clears from the posterior of *lgs* RNAi embryos mainly at 32-35 hours AEL (Figure 7C, D). Hence, the initiation of *Kr* expression in *lgs* RNAi embryos is delayed by 3 hours, while the clearance of *Kr* expression is delayed by 9 hours compared to WT. This indicates a continual dilation of *Kr* regulation, not just a delay in the initiation of its expression. On the other hand, in *nos*;*pum* RNAi embryos, most of the embryos initiated *Kr* expressed at 20-23 hours AEL, manifesting a delay of 3 hours in *Kr* expression initiation compared to WT. *Kr* expression then clears from the posterior end of *nos*;*pum* embryos at 26-29 hours AEL, manifesting a delay of roughly 3 hours compared to WT as well. Hence, *Kr* expression in *nos*;*pum* RNAi embryos suffers a 3 hours delay only during the initiation phase, and this delay is carried over until the end of *Kr* expression without further delay in *Kr* regulation (as clear from Figure 7D).

Same behavior was observed for *mlpt* expression. *mlpt* is expressed in two trunk domains in *Tribolium*. In *lgs* RNAi embryos, a persistent dilation of *mlpt* regulation is observed during both blastoderm and germband stages (Figure 7C, E). In contrast, a delay in the initiation of *mlpt* expression during the blastoderm stage is observed in *nos*;*pum* RNAi embryos that is carried over to the germband stage without further dilation of *mlpt* regulation (Figure 7C, E). These results support our models in which *cad* acts as a speed regulator for gap gene regulation in *Tribolium* while the maternal Hb gradient mediates fast initiation of gap gene expression during the blastoderm stage.

## Conclusion

Recently, we presented a model for gap gene regulation in short- and intermediate-germ insects in which the speed of a genetic cascade is regulated by a posterior-to-anterior gradient of Wnt/Cad (10). The model (termed the ‘Speed Regulation’ model) provided a mechanistic basis for how gap genes are expressed in waves, and how these waves can pattern a non-elongating tissue like the blastoderm as well as an elongating tissue like the germband. This also provided a simple model for the evolution of short-germ insects (in which AP patterning takes place mostly during the germband phase of development) to long-germ insects (in which AP patterning takes place mostly during the blastoderm stage). The model predicted that in long-germ insects, gap genes are expressed in sequential waves that are mediated by a posterior-to-anterior gradient of Wnt/Cad. Although this might be the case in some long-germ insects (possibly long-germ beetles like *Callosobruchus*), it is not the case in other long-germ insects like *Drosophila* (and other Diptera), where gap gene domains arise more or less simultaneously during the blastoderm stage. This simultaneous mode of gene expression potentially evolved to mediate fast AP patterning, as a response to the evolutionary pressure for an overall fast development in *Drosophila*. Furthermore, gap gene regulation in *Drosophila* seems to rely on anterior-to-posterior gradient (namely, those of Bcd and maternal Hb) with seemingly redundant contribution from the posterior-to-anterior morphogen Cad. In this paper, we presented a model for the evolution of a sequential short/intermediate-germ mode of AP patterning to a more simultaneous long-germ mode of patterning. The model is based on the expression of the first gene in the gap gene cascade (*hb*) in an anterior-to-posterior gradient, which results in the differential initialization of the genetic cascade along the AP axis, and consequently, fast simultaneous patterning with slight posterior-to-anterior shift, as observed in *Drosophila*. Besides being a possible evolutionary mechanism, it explains the ability for an anterior-to-posterior gradient like maternal Hb to solely mediate the formation of gap gene expressions along the AP axis of *Drosophila*, potentially with minimal contribution of a posterior-to-anterior morphogen like Cad. The model recapitulates gap gene expression patterns in WT and different manipulations of the maternal Hb gradient in addition to several gap gene mutants.

## Supporting information

## Acknowledgement

This work is supported by an Alexander von Humboldt fellowship to Ezzat El-Sherif.

---

**Movie S1. Effect of changing posterior morphogen concentration and first gene initialization on patterning speed in a molecular realization of the Speed Regulation model.** Simulations shown here correspond to the snap shots shown in Figure 3. The simulations show the behavior of a model of insect embryogenesis under different conditions. **(A)** Low concentration of the speed regulator and the first gene is expressed uniformly along the AP axis (posterior to the right). **(B)** The concentration of the speed regulator is high and the first gene is expressed uniformly along the AP axis. **(C)** The concentration of the speed regulator is *higher* and the first gene is expressed uniformly along the AP axis. **(D)** The concentration of the speed regulator is high and the first gene is expressed in a shallow anterior-to-posterior gradient. **(E)** The concentration of the speed regulator is high and the first gene is expressed in a steep anterior-to-posterior gradient. Simulations backgrounds turn yellow when all gene expression bands form. Speed regulator is shown in grey. Expressions of genes in the genetic cascade (in order): blue, red, green, gold, and brown.

**Movie S2. Module Switching model operating under uniform speed regulator and an anterior-posterior gradient of first gene expression.** Simulation of the Gradual Module Switching model if the speed gradient (grey) is expressed in a posterior-to-anterior gradient, and the first gene in the cascade (blue) is expressed uniformly. Expressions of genes in the genetic cascade (in order): blue, red, green, gold, and brown. The simulation corresponds to the snap shots shown in Figure 4D.

**Movie S3. Model versus Experiment: Effect of varying the dosage of maternal Hb in *Drosophila***. Shown here are simulations of our model of gap gene regulation in *Drosophila* with different dosages of maternal Hb in *bcd*;*tsl* mutant background (**A**: zero dosages, **B**: one dosage, **C**: 4 dosages). Also shown simulation of the model with uniformly expressed maternal Hb (two dosages) in *osk* mutants **(E)**. Simulations shown here correspond to those shown in Figure 5B. Expressions of *hb*, *Kr*, *kni*, and *gt* are shown in blue, red, green, and gold, respectively. Uniform distribution of Cad is shown in grey.

**Movie S4. Model versus Experiment: gap mutants in *Drosophila*.** Simulation of our model for gap gene regulation in *Drosophila* in different genetic background (**A:** WT, **B:** *Kr*-, **C:** *kni*-, **D:** *gt*-). Expressions of *hb*, *Kr*, *kni*, and *gt* are shown in blue, red, green, and gold, respectively. Uniform distribution of Cad is shown in grey.

## Materials and Methods

### *in situ* hybridization, RNAi, and imaging

*In situ* hybridization was performed using DIG-labeled RNA probes and anti-DIG::AP antibody (Roche), and signal was developed using NBT/BCIP (BM Purple, Roche) according to standard protocols (83, 84). All expression analyses were performed using embryos from uninjected females or females injected with double-stranded RNA (dsRNA) of gene of interest. dsRNA was synthesized using the T7 megascript kit (Ambion) and mixed with injection buffer (5 mM KCl, 0.1 mM KPO4, pH 6.8) before injection. Used dsRNA concentrations: 4 µg/µl for both *nos* and *pum* in *nos*;*pum* double RNAi experiments. Embryos were imaged with ProgRes CFcool camera on Zeiss Axio Scope. A1 microscope using ProgRes CapturePro image acquisition software. Brightness and contrast of all images were adjusted and placed on a white background using Adobe Photoshop.

### Egg collections for developmental time windows

Three hours developmental windows were generated by incubating three hours egg collections at 23– 24°C for the desired length of time. Beetles were reared in whole-wheat flour supplemented with 5% dried yeast.

### Calculating temporal profile of gap gene expression at the posterior end of the embryo

For each of the 3-hour developmental windows: 14-17, 17-20, 20-23, 23-26, 26-29, 29-32, 32-35, and 35-38 hours AEL, the percentage of embryos that have detectable expression of the gene of interest (versus embryos with no expression) is calculated and used as an estimate for the gene expression level at that point of time. Figures 7D, E are produced using this methodology.

### Computational Modeling

See Text S1.

## References

1. Palmeirim I, Henrique D, Ish-Horowicz D, Pourquié O. Avian hairy gene expression identifies a molecular clock linked to vertebrate segmentation and somitogenesis. Cell. 1997 Nov 28;91(5):639–648.

2. Dequéant ML, Glynn E, Gaudenz K, Wahl M, Chen J, Mushegian A, et al. A complex oscillating network of signaling genes underlies the mouse segmentation clock. Science. 2006 Dec 8;314(5805):1595–1598.

3. Li Y, Fenger U, Niehrs C, Pollet N. Cyclic expression of esr9 gene in Xenopus presomitic mesoderm. Differentiation. 2003 Jan 1;71(1):83–89.

4. Moreno-Risueno MA, Van Norman JM, Moreno A, Zhang J, Ahnert SE, Benfey PN. Oscillating gene expression determines competence for periodic Arabidopsis root branching. Science. 2010 Sep 10;329(5997):1306–1311.

5. Petricka JJ, Winter CM, Benfey PN. Control of Arabidopsis root development. Annu Rev Plant Biol. 2012 Feb 9;63:563–590.

6. Kohwi M, Doe CQ. Temporal fate specification and neural progenitor competence during development. Nat Rev Neurosci. 2013 Nov 20;14(12):823–838.

7. Li X, Erclik T, Bertet C, Chen Z, Voutev R, Venkatesh S, et al. Temporal patterning of Drosophila medulla neuroblasts controls neural fates. Nature. 2013 Jun 27;498(7455):456–462.

8. Dessaud E, Yang LL, Hill K, Cox B, Ulloa F, Ribeiro A, et al. Interpretation of the sonic hedgehog morphogen gradient by a temporal adaptation mechanism. Nature. 2007 Nov 29;450(7170):717–720.

9. Deschamps J, Duboule D. Embryonic timing, axial stem cells, chromatin dynamics, and the Hox clock. Genes Dev. 2017 Jul 15;31(14):1406–1416.

10. Zhu X, Rudolf H, Healey L, François P, Brown SJ, Klingler M, et al. Speed regulation of genetic cascades allows for evolvability in the body plan specification of insects. Proc Natl Acad Sci U S A. 2017 Sep 25;114(41).

11. Clark E, Peel AD. Evidence for the temporal regulation of insect segmentation by a conserved sequence of transcription factors. Development. 2018 May 3

12. Lauschke VM, Tsiairis CD, François P, Aulehla A. Scaling of embryonic patterning based on phase-gradient encoding. Nature. 2013 Jan 3;493(7430):101–105.

13. Oates AC, Morelli LG, Ares S. Patterning embryos with oscillations: structure, function and dynamics of the vertebrate segmentation clock. Development. 2012 Feb;139(4):625–639.

14. Dubrulle J, McGrew MJ, Pourquié O. tFGF signaling controls somite boundary position and regulates segmentation clock control of spatiotemporal Hox gene activation. Cell. 2001 Jul 27;106(2):219–232.

15. Sonnen KF, Lauschke VM, Uraji J, Falk HJ, Petersen Y, Funk MC, et al. tModulation of Phase Shift between Wnt and Notch Signaling Oscillations Controls Mesoderm Segmentation. Cell. 2018 Feb 22;172(5):1079–1090.e12.

16. Deschamps J, van Nes J. Developmental regulation of the Hox genes during axial morphogenesis in the mouse. Development. 2005 Jul 1;132(13):2931–2942.

17. Gaunt SJ. Gradients and forward spreading of vertebrate Hox gene expression detected by using a Hox/lacZ transgene. Dev Dyn. 2001 May 1;221(1):26–36.

18. Balaskas N, Ribeiro A, Panovska J, Dessaud E, Sasai N, Page KM, et al. Gene regulatory logic for reading the Sonic Hedgehog signaling gradient in the vertebrate neural tube. Cell. 2012 Jan 20;148(1-2):273–284.

19. Cohen M, Page KM, Perez-Carrasco R, Barnes CP, Briscoe J. A theoretical framework for the regulation of Shh morphogen-controlled gene expression. Development. 2014 Oct;141(20):3868–3878.

20. Briscoe J, Small S. Morphogen rules: design principles of gradient-mediated embryo patterning. Development. 2015 Dec 1;142(23):3996–4009.

21. Uzkudun M, Marcon L, Sharpe J. Data-driven modelling of a gene regulatory network for cell fate decisions in the growing limb bud. Mol Syst Biol. 2015 Jul 14;11(7):815.

22. Towers M, Tickle C. Growing models of vertebrate limb development. Development. 2009 Jan;136(2):179–190.

23. Summerbell D, Lewis JH, Wolpert L. Positional information in chick limb morphogenesis. Nature. 1973 Aug 24;244(5417):492–496.

24. Lynch JA, El-Sherif E, Brown SJ. Comparisons of the embryonic development of Drosophila, Nasonia, and Tribolium. Wiley Interdiscip Rev Dev Biol. 2012 Feb;1(1):16–39.

25. Jaeger J. The gap gene network. Cell Mol Life Sci. 2011 Jan;68(2):243–274.

26. Jaeger J, Surkova S, Blagov M, Janssens H, Kosman D, Kozlov KN, et al. Dynamic control of positional information in the early Drosophila embryo. Nature. 2004 Jul 15;430(6997):368–371.

27. El-Sherif E, Levine M. Shadow enhancers mediate dynamic shifts of gap gene expression in the drosophila embryo. Curr Biol. 2016 May 9;26(9):1164–1169.

28. Lim B, Fukaya T, Heist T, Levine M. Temporal dynamics of pair-rule stripes in living Drosophila embryos. Proc Natl Acad Sci U S A. 2018 Aug 14;115(33):8376–8381.

29. Berrocal A, Lammers NC, Garcia HG, Eisen MB. Kinetic sculpting of the seven stripes of the Drosophila even-skipped gene. BioRxiv. 2018 May 31;

30. El-Sherif E, Zhu X, Fu J, Brown SJ. Caudal regulates the spatiotemporal dynamics of pair-rule waves in Tribolium. PLOS Genet. 2014 Oct 16;10(10):e1004677.

31. El-Sherif E, Averof M, Brown SJ. A segmentation clock operating in blastoderm and germband stages of Tribolium development. Development. 2012 Dec 1;139(23):4341–4346.

32. Davis GK, Patel NH. Short, long, and beyond: molecular and embryological approaches to insect segmentation. annu Rev Entomol. 2002;47:669–699.

33. Patel NH, Hayward DC, Lall S, Pirkl NR, DiPietro D, Ball EE. Grasshopper hunchback expression reveals conserved and novel aspects of axis formation and segmentation. Development. 2001 Sep;128(18):3459–3472.

34. Rosenberg MI, Brent AE, Payre F, Desplan C. Dual mode of embryonic development is highlighted by expression and function of Nasonia pair-rule genes. elife. 2014 Mar 5;3(3):e01440.

35. Patel NH, Condron BG, Zinn K. Pair-rule expression patterns of even-skipped are found in both short-and long-germ beetles. Nature. 1994 Feb 3;367(6462):429–434.

36. Stahi R, Chipman AD. Blastoderm segmentation in Oncopeltus fasciatus and the evolution of insect segmentation mechanisms. Proc Biol Sci. 2016 Oct 12;283(1840).

37. Auman T, Chipman AD. Growth zone segmentation in the milkweed bug Oncopeltus fasciatus sheds light on the evolution of insect segmentation. BioRxiv. 2018 May 21;

38. Xiang J, Forrest IS, Pick L. Dermestes maculatus: an intermediate-germ beetle model system for evo-devo. Evodevo. 2015 Oct 16;6(1):32.

39. Sucena É, Vanderberghe K, Zhurov V, Grbić M. Reversion of developmental mode in insects: evolution from long germband to short germband in the polyembrionic wasp Macrocentrus cingulum Brischke. Evol Dev. 2014 Aug;16(4):233–246.

40. Kuhlmann L, El-Sherif E. Speed regulation and gradual enhancer switching models as flexible and evolvable patterning mechanisms. BioRxiv. 2018 Feb 7;

41. Murray JD. Mathematical Biology: I. Jan Introduction (Interdisciplinary Applied Mathematics) (Pt. 1) [Internet]. 3rd ed. Springer; 2007. Available from: https://www.amazon.com/Mathematical-Mathematics/dp/0387952233Biology-Introduction-Interdisciplinary-Mathematics/dp/0387952233

42. Winfree AT. The Geometry of Biological Time (Interdisciplinary Applied Mathematics) [Internet]. 2nd ed. 2001. Softcover reprint of the original 2nd ed. 2001. Springer; 2010. Available from: https://www.amazon.com/Geometry-Biological-Interdisciplinary-Applied-Mathematics/dp/1441931961

43. beck MT, Váradi ZB. One, Two and Three-dimensional Spatially Periodic Chemical Reactions. Nature [Internet]. 1972 Jan 3; Available from: http://www.nature.com/nature-physci/journal/v235/n53/abs/physci235015a0.html

44. thoenes D. Spatial Oscillations” in the Zhabotinskii Reaction. Nature [Internet]. 1973 May 14; Available from: http://www.nature.com/nature-physci/journal/v243/n124/abs/physci243018a0.html

45. Boos A, Distler J, Rudolf H, Klingler M, El-Sherif E. A re-inducible genetic cascade patterns the anterior-posterior axis of insects in a threshold-free fashion. BioRxiv. 2018 May 15;

46. Olesnicky EC, Brent AE, Tonnes L, Walker M, Pultz MA, Leaf D, et al. A caudal mRNA gradient controls posterior development in the wasp Nasonia. Development. 2006 Oct;133(20):3973–3982.

47. Verd B, Clark E, Wotton KR, Janssens H, Jiménez-Guri E, Crombach A, et al. A damped oscillator imposes temporal order on posterior gap gene expression in Drosophila. PLOS Biol. 2018 mFeb 16;16(2):e2003174.

48. Stauber M, Prell A, Schmidt-Ott U. A single Hox3 gene with composite bicoid and zerknullt expression characteristics in non-Cyclorrhaphan flies. Proc Natl Acad Sci U S A. 2002 Jan 8;99(1):274–279.

49. Brown S, Fellers J, Shippy T, Denell R, Stauber M, Schmidt-Ott U. A strategy for mapping bicoid on the phylogenetic tree. Curr Biol. 2001 Jan 23;11(2):R43–4.

50. Falciani F, Hausdorf B, Schröder R, Akam M, Tautz D, Denell R, et al. Class 3 Hox genes in insects and the origin of zen. Proc Natl Acad Sci U S A. 1996 Aug 6;93(16):8479–8484.

51. Hülskamp M, Pfeifle C, Tautz D. A morphogenetic gradient of hunchback protein organizes the expression of the gap genes Krüppel and knirps in the early Drosophila embryo. Nature. 1990 Aug 9;346(6284):577–580.

52. Tufcea DE, François P. Critical Timing without a Timer for Embryonic Development. Biophys J. 2015 Oct 20;109(8):1724–1734.

53. Hastings A, Abbott KC, Cuddington K, Francis T, Gellner G, Lai YC, et al. Transient phenomena in ecology. Science. 2018 Sep 7;361(6406).

54. Driever W, Nüsslein-Volhard C. The bicoid protein determines position in the Drosophila embryo in a concentration-dependent manner. Cell. 1988 Jul 1;54(1):95–104.

55. Struhl G, Struhl K, Macdonald PM. The gradient morphogen bicoid is a concentration-dependent transcriptional activator. Cell. 1989 Jun 30;57(7):1259–1273.

56. Struhl G, Johnston P, Lawrence PA. Control of Drosophila body pattern by the hunchback morphogen gradient. Cell. 1992 Apr 17;69(2):237–249.

57. Rivera-Pomar R, Lu X, Perrimon N, Taubert H, Jäckle H. Activation of posterior gap gene expression in the Drosophila blastoderm. Nature. 1995 Jul 20;376(6537):253–256.

58. Curtis D, Apfeld J, Lehmann R. nanos is an evolutionarily conserved organizer of anterior-posterior polarity. Development. 1995 Jun;121(6):1899–1910.

59. Barker DD, Wang C, Moore J, Dickinson LK, Lehmann R. Pumilio is essential for function but not for distribution of the Drosophila abdominal determinant Nanos. Genes Dev. 1992 Dec;6(12A):2312–2326.

60. Rivera-Pomar R, Niessing D, Schmidt-Ott U, Gehring WJ, Jäckle H. RNA binding and translational suppression by bicoid. Nature. 1996 mFeb 22;379(6567):746–749.

61. Schulz C, Tautz D. Zygotic caudal regulation by hunchback and its role in abdominal segment formation of the Drosophila embryo. Development. 1995 Apr;121(4):1023–1028.

62. Wreden C, Verrotti AC, Schisa JA, Lieberfarb ME, Strickland S. Nanos and pumilio establish embryonic polarity in Drosophila by promoting posterior deadenylation of hunchback mRNA. Development. 1997 Aug;124(15):3015–3023.

63. Yu D, Small S. Precise registration of gene expression boundaries by a repressive morphogen in Drosophila. Curr Biol. 2008 Jun 24;18(12):868–876.

64. Lehmann R, Nüsslein-Volhard C. The maternal gene nanos has a central role in posterior pattern formation of the Drosophila embryo. Development. 1991 Jul;112(3):679–691.

65. Rothe M, Wimmer EA, Pankratz MJ, González-Gaitán M, Jäckle H. Identical transacting factor requirement for knirps and knirps-related Gene expression in the anterior but not in the posterior region of the Drosophila embryo. Mech Dev. 1994 Jun;46(3):169–181.

66. Pankratz MJ, Hoch M, Seifert E, Jäckle H. Krüppel requirement for knirps enhancement reflects overlapping gap gene activities in the Drosophila embryo. Nature. 1989 Sep 28;341(6240):337–340.

67. Jäckle H, Tautz D, Schuh R, Seifert E, Lehmann R. Cross-regulatory interactions among the gap genes of Drosophila. Nature. 1986 Dec;324(6098):668–670.

68. Hülskamp M, Lukowitz W, Beermann A, Glaser G, Tautz D. Differential regulation of target genes by different alleles of the segmentation gene hunchback in Drosophila. Genetics. 1994 Sep;138(1):125–134.

69. Clyde DE, Corado MS, Wu X, Paré A, Papatsenko D, Small S. A self-organizing system of repressor gradients establishes segmental complexity in Drosophila. Nature. 2003 Dec 18;426(6968):849–853.

70. Kraut R, Levine M. Spatial regulation of the gap gene giant during Drosophila development. Development. 1991 Feb;111(2):601–609.

71. Capovilla M, Eldon ED, Pirrotta V. The giant gene of Drosophila encodes a b-ZIP DNA-binding protein that regulates the expression of other segmentation gap genes. Development. 1992 Jan;114(1):99–112.

72. Eldon ED, Pirrotta V. Interactions of the Drosophila gap gene giant with maternal and zygotic pattern-forming genes. Development. 1991 Feb;111(2):367–378.

73. Schmitt-Engel C, Cerny AC, Schoppmeier M. A dual role for nanos and pumilio in anterior and posterior blastodermal patterning of the short-germ beetle Tribolium castaneum. Dev Biol. 2012 Apr 15;364(2):224–235.

74. Wolff C, Schröder R, Schulz C, Tautz D, Klingler M. Regulation of the Tribolium homologues of caudal and hunchback in Drosophila: evidence for maternal gradient systems in a short germ embryo. Development. 1998 Sep;125(18):3645–3654.

75. Marques-Souza H, Aranda M, Tautz D. Delimiting the conserved features of hunchback function for the trunk organization of insects. Development. 2008 Mar;135(5):881–888.

76. Cerny AC, Bucher G, Schröder R, Klingler M. Breakdown of abdominal patterning in the Tribolium Kruppel mutant jaws. Development. 2005 Dec;132(24):5353–5363.

77. Savard J, Marques-Souza H, Aranda M, Tautz D. A segmentation gene in tribolium produces a polycistronic mRNA that codes for multiple conserved peptides. Cell. 2006 Aug 11;126(3):559–569.

78. Bucher G, Klingler M. Divergent segmentation mechanism in the short germ insect Tribolium revealed by giant expression and function. Development. 2004 Apr;131(8):1729–1740.

79. Marques-Souza H. Evolution of the gene regulatory network controlling trunk segmentation in insects [Internet] [Doctoral dissertation]. University of Cologne; 2007. Available from: http://kups.ub.uni-koeln.de/2167/

80. Bolognesi R, Beermann A, Farzana L, Wittkopp N, Lutz R, Balavoine G, et al. Tribolium Wnts: evidence for a larger repertoire in insects with overlapping expression patterns that suggest multiple redundant functions in embryogenesis. Dev Genes Evol. 2008 Apr 8;218(3-4):193–202.

81. Bolognesi R, Farzana L, Fischer TD, Brown SJ. Multiple Wnt genes are required for segmentation in the short-germ embryo of Tribolium castaneum. Curr Biol. 2008 Oct 28;18(20):1624–1629.

82. Fu J, Posnien N, Bolognesi R, Fischer TD, Rayl P, Oberhofer G, et al. Asymmetrically expressed axin required for anterior development in Tribolium. Proc Natl Acad Sci U S A. 2012 May 15;109(20):7782–7786.

83. Schinko J, Posnien N, Kittelmann S, Koniszewski N, Bucher G. Single and double whole-mount in situ hybridization in red flour beetle (Tribolium) embryos. Cold Spring Harb Protoc. 2009 Aug;2009(8):pdb.prot5258.

84. Shippy TD, Coleman CM, Tomoyasu Y, Brown SJ. Concurrent in situ hybridization and antibody staining in red flour beetle (Tribolium) embryos. Cold Spring Harb Protoc. 2009 Aug;2009(8):pdb.prot5257.

