## Supplementary material for "Speeding up anterior-posterior patterning of insects by differential initialization of the gap gene cascade"

### Supplementary Text S1. Computational Modeling

All computational models were created using Matlab/Octave. For all GRN models in this study (except for AC-DC motif; see below), the activity of an enhancer ***E*** of gene ***X*** to which $N_a$ activators ($\boldsymbol{A}_i$, $i$=1 to $N_a$) and $N_r$ repressors ($\boldsymbol{R}_j$, $j$=1 to $N_r$) bind is modelled using AND gated Hill functions of individual binding proteins:

$$E = \prod_{i=1}^{Na} \frac{(A_i/K_i)^{na_i}}{1 + (A_i/K_i)^{na_i}} \prod_{j=1}^{Nr} \frac{1}{1 + (R_j/K_j')^{nr_j}}$$

where $E$ is the activity of enhancer ***E***, $A_i$ is the concentration of activator $\boldsymbol{A}_i$, $R_j$ is the concentration of repressor $\boldsymbol{R}_j$, $na_i$ is the cooperativity of $\boldsymbol{A}_i$ binding to the enhancer **E**, $nr_j$ is the cooperativity of $\boldsymbol{R}_j$ binding to the enhancer ***E***, $K_i$ is the dissociation constant of activator $\boldsymbol{A}_i$, and $K'_j$ is the dissociation constant of repressor $\boldsymbol{R}_j$.

### Speed Regulation GRN (Module Switching model)

The transcription rate of gene ***X*** ($X$) regulated by two separate modules ($\boldsymbol{E}_1$ and $\boldsymbol{E}_2$) is modeled as a weighted sum of the activity of the two modules: $= c_1E_1 + c_2E_2$ . The following are the equations used for modeling the 5-genes gradual module switching model. $X_{Di}$ is the activity of the dynamic module of gene $\boldsymbol{X}_i$. $X_{Si}$ is the activity of the static module of gene $\boldsymbol{X}_i$. $\boldsymbol{X}_i$ is the mRNA concentration transcribed by gene $\boldsymbol{X}_i$. $G$ is the concentration of the (speed regulator) gradient ***G***.

<u>Dynamic Modules</u>:

$$\frac{dX_{D1}}{dt} = \frac{G}{1+G} \times \frac{1}{1+\left(\frac{X_2}{0.4}\right)^5} \times \frac{1}{1+\left(\frac{X_3}{0.4}\right)^5} \times \frac{1}{1+\left(\frac{X_4}{0.4}\right)^5} \times \frac{1}{1+\left(\frac{X_5}{0.4}\right)^5}$$

$$\frac{dX_{D2}}{dt} = \frac{G}{1+G} \times \frac{1}{1+\left(\frac{X_1}{2.5}\right)^5} \times \frac{1}{1+\left(\frac{X_3}{0.4}\right)^5} \times \frac{1}{1+\left(\frac{X_3}{0.4}\right)^5} \times \frac{1}{1+\left(\frac{X_5}{0.4}\right)^5}$$

$$\frac{dX_{D3}}{dt} = \frac{G}{1+G} \times \frac{1}{1+\left(\frac{X_1}{0.4}\right)^5} \times \frac{1}{1+\left(\frac{X_2}{2.5}\right)^5} \times \frac{1}{1+\left(\frac{X_4}{0.4}\right)^5} \times \frac{1}{1+\left(\frac{X_5}{0.4}\right)^5}$$

$$\frac{dX_{D4}}{dt} = \frac{G}{1+G} \times \frac{1}{1+\left(\frac{X_1}{0.4}\right)^5} \times \frac{1}{1+\left(\frac{X_2}{0.4}\right)^5} \times \frac{1}{1+\left(\frac{X_3}{2.5}\right)^5} \times \frac{1}{1+\left(\frac{X_5}{0.4}\right)^5}$$

$$\frac{dX_{D5}}{dt} = \frac{G}{1+G} \times \frac{1}{1+\left(\frac{X_1}{0.4}\right)^5} \times \frac{1}{1+\left(\frac{X_2}{0.4}\right)^5} \times \frac{1}{1+\left(\frac{X_3}{0.4}\right)^5} \times \frac{1}{1+\left(\frac{X_4}{2.5}\right)^5}$$

Static Modules:

$$\frac{dX_{S1}}{dt} = \frac{1}{1+G} \times \frac{1}{1+\left(\frac{X_2}{0.4}\right)^5} \times \frac{1}{1+\left(\frac{X_3}{0.4}\right)^5} \times \frac{1}{1+\left(\frac{X_4}{0.4}\right)^5} \times \frac{1}{1+\left(\frac{X_5}{0.4}\right)^5}$$

$$\frac{dX_{S2}}{dt} = \frac{1}{1+G} \times \frac{1}{1+\left(\frac{X_1}{0.4}\right)^5} \times \frac{1}{1+\left(\frac{X_3}{0.4}\right)^5} \times \frac{1}{1+\left(\frac{X_4}{0.4}\right)^5} \times \frac{1}{1+\left(\frac{X_5}{0.4}\right)^5}$$

$$\frac{dX_{S3}}{dt} = \frac{1}{1+G} \times \frac{1}{1+\left(\frac{X_1}{0.4}\right)^5} \times \frac{1}{1+\left(\frac{X_2}{0.4}\right)^5} \times \frac{1}{1+\left(\frac{X_4}{0.4}\right)^5} \times \frac{1}{1+\left(\frac{X_5}{0.4}\right)^5}$$

$$\frac{dX_{S4}}{dt} = \frac{1}{1+G} \times \frac{1}{1+\left(\frac{X_1}{0.4}\right)^5} \times \frac{1}{1+\left(\frac{X_2}{0.4}\right)^5} \times \frac{1}{1+\left(\frac{X_3}{0.4}\right)^5} \times \frac{1}{1+\left(\frac{X_5}{0.4}\right)^5}$$

$$\frac{dX_{S5}}{dt} = \frac{1}{1+G} \times \frac{1}{1+\left(\frac{X_1}{0.4}\right)^5} \times \frac{1}{1+\left(\frac{X_2}{0.4}\right)^5} \times \frac{1}{1+\left(\frac{X_3}{0.4}\right)^5} \times \frac{1}{1+\left(\frac{X_4}{0.4}\right)^5}$$

Combining the activities of Dynamic and Static Enhancers:

$$\frac{dX_1}{dt} = 3\frac{dX_{D1}}{dt} + 2\frac{dX_{S1}}{dt} - X_1$$

$$\frac{dX_2}{dt} = 3\frac{dX_{D2}}{dt} + 2\frac{dX_{S2}}{dt} - X_2$$

$$\frac{dX_3}{dt} = 3\frac{dX_{D3}}{dt} + 2\frac{dX_{S3}}{dt} - X_3$$

$$\frac{dX_4}{dt} = 3\frac{dX_{D4}}{dt} + 2\frac{dX_{S4}}{dt} - X_4$$

$$\frac{dX_5}{dt} = 3\frac{dX_{D5}}{dt} + 2\frac{dX_{S5}}{dt} - X_5$$

All simulations in this study are based on this model, with different number of genes, gradient

(***G***) dynamics, and initial conditions. Below, we provide the Matlab/Octave codes corresponding to each simulation in each Supplementary Movie.

Matlab/Octave code for Movie S1

```
function MovieS1
clear
clc
close all

%Enter which simulation option you would like to run: A, B, C, D, or E
%These correspond to simulations in Figure 3 A, B, C, D, and E
%and Movie S1 A, B, C, D, and E
option= 'E';

%Settign parameters that differ between Movies S1: A, B, C, D, and E
% (1) AP axis length differs in each case to accommodate all gene expression
% bands .. adjustable parameter: AP_axis_length
% (2) The level of the posterior gradient (speed regulator)... adjustable
% parameter: speed_gradient_level
% (3) Expression of the first gene: exp(-alpha*AP_axis)
if option=='A'
    AP_axis_length=0.65;
    speed_gradient_level=0.3;
    alpha=0;
elseif option=='B'
    AP_axis_length=0.4;
    speed_gradient_level=1.5;
    alpha=0;
elseif option=='C'
    AP_axis_length=0.4;
    speed_gradient_level=2.5;
    alpha=0;
elseif option=='D'
    AP_axis_length=0.25;
    speed_gradient_level=1.5;
    alpha=10;
elseif option=='E'
    AP_axis_length=0.15;
    speed_gradient_level=1.5;
    alpha=35;
else
    error('Available options are either A, B, C, D, or E')
end

%Time Axis
t=0:.01:15;

% AP_axis
AP_axis=0:.001:AP_axis_length;

%Posterior (speed regulator) gradient parameters
b_to_g_time=2;%time of blastoderm to germband transition
flat_start=.1;%position of start of flate region of the gradient in the
blastoderm
v=.05;%wavefront velocity during germband stage

%solve the model
```

```
spacetime=zeros(5,length(t),length(AP_axis));
for pos=1:length(AP_axis)
    %initial conditions for genes 1, 2, 3, 4, 5 .. in addition parameters
    %passed to the differential equations
    initial_conditions=[exp(-alpha*AP_axis(pos)) 0 0 0 0 AP_axis(pos)
b_to_g_time flat_start v speed_gradient_level];
    [T, x]= ode45(@odefun,t,initial_conditions);
    spacetime(:,:,pos)=x(:,1:5)';
end

%plot solution
close all
for nt=1:length(t)

    %posterior gradient (speed regulator) set up
    if(t(nt)<b_to_g_time)

g=speed_gradient_level*(AP_axis/flat_start).^3./(1+(AP_axis/flat_start).^3);
    else
        n=3*exp(t(nt)-b_to_g_time);
        if(n>100)
            n=100;
        end
        g=speed_gradient_level*(AP_axis/(flat_start+v*(t(nt)-
b_to_g_time))).^n./(1+(AP_axis/(flat_start+v*(t(nt)-b_to_g_time))).^n);
    end

    set(gca, 'ColorOrder', [86 128 193; 229 51 50; 79 185 118; 219 152 40;
168 124 78;128 128 128 ]/256, 'NextPlot', 'replacechildren');
    plot(AP_axis,[squeeze(spacetime(:,nt,:))' g'],'LineWidth',5)
    axis([0 max(AP_axis) 0 2.5])

    set(gca,'xtick',[])
    set(gca,'ytick',[])
    pause(.00000001)
end

%%%%% Model %%%%%%
function dx = odefun(t,x)

x1=x(1);
x2=x(2);
x3=x(3);
x4=x(4);
x5=x(5);
AP_axis=x(6);
b_to_g_time=x(7);
flat_start=x(8);
v=x(9);
speed_gradient_level=x(10);

%posterior gradient (speed regulator) set up
if(t<b_to_g_time)
    g=speed_gradient_level*(AP_axis/flat_start)^3/(1+(AP_axis/flat_start)^3);
else
```

```
    n=(3*exp(t-b_to_g_time));
    if(n>100)
        n=100;
    end
    g=speed_gradient_level*(AP_axis/(flat_start+v*(t-
b_to_g_time)))^n/(1+(AP_axis/(flat_start+v*(t-b_to_g_time)))^n);
end

w=2.5;%Dissociation constant for weak repressions
s=.4;%Dissociation constant for strong repressions
n=5;%cooperativity
lambda=1;%mRNA decay rate

%dynamic module
dynamic(1)=
g/(1+g)*1/(1+(x2/s)^n)*1/(1+(x3/s)^n)*1/(1+(x4/s)^n)*1/(1+(x5/s)^n);
dynamic(2)=
g/(1+g)*1/(1+(x1/w)^n)*1/(1+(x3/s)^n)*1/(1+(x4/s)^n)*1/(1+(x5/s)^n);
dynamic(3)=
g/(1+g)*1/(1+(x1/s)^n)*1/(1+(x2/w)^n)*1/(1+(x4/s)^n)*1/(1+(x5/s)^n);
dynamic(4)=
g/(1+g)*1/(1+(x1/s)^n)*1/(1+(x2/s)^n)*1/(1+(x3/w)^n)*1/(1+(x5/s)^n);
dynamic(5)=
g/(1+g)*1/(1+(x1/s)^n)*1/(1+(x2/s)^n)*1/(1+(x3/s)^n)*1/(1+(x4/w)^n);

% static module
static(1)=
(1/(g+1))*1/(1+(x2/s)^n)*1/(1+(x3/s)^n)*1/(1+(x4/s)^n)*1/(1+(x5/s)^n);
static(2)=
(1/(g+1))*1/(1+(x1/s)^n)*1/(1+(x3/s)^n)*1/(1+(x4/s)^n)*1/(1+(x5/s)^n);
static(3)=
(1/(g+1))*1/(1+(x1/s)^n)*1/(1+(x2/s)^n)*1/(1+(x4/s)^n)*1/(1+(x5/s)^n);
static(4)=
(1/(g+1))*1/(1+(x1/s)^n)*1/(1+(x2/s)^n)*1/(1+(x3/s)^n)*1/(1+(x5/s)^n);
static(5)=
(1/(g+1))*1/(1+(x1/s)^n)*1/(1+(x2/s)^n)*1/(1+(x3/s)^n)*1/(1+(x4/s)^n);

%total gene regulation: dynamic module contribution + static module
%contribution - decay
c=2;
d=3;
dx(1)= c*static(1)+d*dynamic(1)-lambda*x1;
dx(2)= c*static(2)+d*dynamic(2)-lambda*x2;
dx(3)= c*static(3)+d*dynamic(3)-lambda*x3;
dx(4)= c*static(4)+d*dynamic(4)-lambda*x4;
dx(5)= c*static(5)+d*dynamic(5)-lambda*x5;

%zero change in parameters
dx(6)=0;
dx(7)=0;
dx(8)=0;
dx(9)=0;
dx(10)=0;

dx=dx';
```

Matlab/Octave code for Movie S2

```
function MovieS2

clear
clc
close all

%Time Axis
t=0:.01:6;

% AP_axis
AP_axis=0:.001:.15;

%solve the model
spacetime=zeros(5,length(t),length(AP_axis));
for pos=1:length(AP_axis)
    %initial conditions for genes 1, 2, 3, 4, 5 .. in addition parameters
    %passed to the differential equations
    initial_conditions=[exp(-35*AP_axis(pos)) 0 0 0 0 length(AP_axis)];
    [T, x]= ode45(@odefun,t,initial_conditions);
    spacetime(:,:,pos)=x(:,1:5)';
end

%plot solution
close all
for nt=1:length(t)

    %posterior gradient (speed regulator) set up
    if (t(nt)<=2)
        g= ones(1,151)-0.5-t(nt)/4;
    else
        g=zeros(1,151);
    end

    set(gca, 'ColorOrder', [86 128 193; 229 51 50; 79 185 118; 219 152 40;
168 124 78;128 128 128 ]/256, 'NextPlot', 'replacechildren');

    plot(AP_axis,[squeeze(spacetime(:,nt,:))' g'],'LineWidth',5)
    axis([0 max(AP_axis) 0 max(max(max(spacetime)))])

    axis off
    pause(.00000001)
end

%%%%% Model %%%%%%
function dx = odefun(t,x)

x1=x(1);
x2=x(2);
x3=x(3);
x4=x(4);
x5=x(5);
AP_axis_length=x(6);
```

```
%posterior gradient (speed regulator) set up
if (t<=2)
    g= ones(1,AP_axis_length)-0.5 - t/4;
else
    g=zeros(1,AP_axis_length);
end

w=2.5;%Dissociation constant for weak repressions
s=.4;%Dissociation constant for strong repressions
n=5;%cooperativity
lambda=1;%mRNA decay rate

% dynamic module
dynamic(1)=
g/(1+g)*1/(1+(x2/s)^n)*1/(1+(x3/s)^n)*1/(1+(x4/s)^n)*1/(1+(x5/s)^n);
dynamic(2)=
g/(1+g)*1/(1+(x1/w)^n)*1/(1+(x3/s)^n)*1/(1+(x4/s)^n)*1/(1+(x5/s)^n);
dynamic(3)=
g/(1+g)*1/(1+(x1/s)^n)*1/(1+(x2/w)^n)*1/(1+(x4/s)^n)*1/(1+(x5/s)^n);
dynamic(4)=
g/(1+g)*1/(1+(x1/s)^n)*1/(1+(x2/s)^n)*1/(1+(x3/w)^n)*1/(1+(x5/s)^n);
dynamic(5)=
g/(1+g)*1/(1+(x1/s)^n)*1/(1+(x2/s)^n)*1/(1+(x3/s)^n)*1/(1+(x4/w)^n);

% static module
static(1)=
ones(1,AP_axis_length)/(1+g)*1/(1+(x2/s)^n)*1/(1+(x3/s)^n)*1/(1+(x4/s)^n)*1/(
1+(x5/s)^n);
static(2)=
ones(1,AP_axis_length)/(1+g)*1/(1+(x1/s)^n)*1/(1+(x3/s)^n)*1/(1+(x4/s)^n)*1/(
1+(x5/s)^n);
static(3)=
ones(1,AP_axis_length)/(1+g)*1/(1+(x1/s)^n)*1/(1+(x2/s)^n)*1/(1+(x4/s)^n)*1/(
1+(x5/s)^n);
static(4)=
ones(1,AP_axis_length)/(1+g)*1/(1+(x1/s)^n)*1/(1+(x2/s)^n)*1/(1+(x3/s)^n)*1/(
1+(x5/s)^n);
static(5)=
ones(1,AP_axis_length)/(1+g)*1/(1+(x1/s)^n)*1/(1+(x2/s)^n)*1/(1+(x3/s)^n)*1/(
1+(x4/s)^n);

%total gene regulation: dynamic module contribution + static module
%contribution - decay
c=2;
d=3;
dx(1)= c*static(1)+d*dynamic(1)-lambda*x1;
dx(2)= c*static(2)+d*dynamic(2)-lambda*x2;
dx(3)= c*static(3)+d*dynamic(3)-lambda*x3;
dx(4)= c*static(4)+d*dynamic(4)-lambda*x4;
dx(5)= c*static(5)+d*dynamic(5)-lambda*x5;

%zero change in parameters
dx(6)=0;
dx=dx';
```

Matlab/Octave code for Movie S3

```
function MovieS3

clear
clc
close all

%Enter which simulation option you would like to run: A, B, C, D, or E
%These correspond to simulations in Figure 5: Hb dosage 0x, 1x, 2x, 4x,
%osk-
%and Movie S3 A, B, C, D, and E
option= 'A';

%Settign parameters that differ between Movies S1: A, B, C, D, and E
% (1) uniform: equals 1 if the maternal Hb is uniform along the AP axis,
% and 0 otherwise
% (2) maternal_Hb_dosage

if option=='A' % 0x maternal Hb dosage
    uniform=0;
    maternal_Hb_dosage=0;
elseif option=='B' % 1x maternal Hb dosage
    uniform=0;
    maternal_Hb_dosage=1;
elseif option=='C' % 2x maternal Hb dosage
    uniform=0;
    maternal_Hb_dosage=2;
elseif option=='D' % 4x maternal Hb dosage
    uniform=0;
    maternal_Hb_dosage=4;
elseif option=='E' % osk- (uniform maternal Hb gradient)
    uniform=1;
    maternal_Hb_dosage=2;
else
    error('Available options are either A, B, C, D, or E')
end

%Time Axis
t=0:.01:4;

% AP_axis
AP_axis=0:.001:.6;

%solve model
spacetime=zeros(4,length(t),length(AP_axis));
for pos=1:length(AP_axis)
    %initial conditions for Drosophila genes hb, Kr, kni, gt
    initial_conditions=[maternal_Hb_dosage*uniform+(1-
uniform)*maternal_Hb_dosage*exp(-8*AP_axis(pos)) 0 0 0];
    [T, x]= ode45(@odefun,t,initial_conditions);
    spacetime(:,:,pos)=x(:,1:4)';
end

%plot solution
```

```
close all
for nt=1:length(t)

    %posterior gradient (speed regulator) set up
    g=0.3*exp(-0.2*t(nt))*ones(size(AP_axis));

    set(gca, 'ColorOrder', [86 128 193; 229 51 50; 79 185 118; 219 152 40;
128 128 128 ]/256, 'NextPlot', 'replacechildren');

    plot(AP_axis,[squeeze(spacetime(:,nt,:))' g'],'LineWidth',5)
    axis([0 max(AP_axis) 0 max(max(max(spacetime)))])

    axis off
    pause(.00000001)
end

%%%%% Model %%%%%%
function dx = odefun(t,x)
x1=x(1);%hb
x2=x(2);%Kr
x3=x(3);%kni
x4=x(4);%gt

w=2.5;%Dissociation constant for weak repressions
s=.4;%Dissociation constant for strong repressions
lambda=1;%mRNA decay rate

%posterior gradient (speed regulator) set up
g=0.3*exp(-0.2*t);

%dynamic module
n=5;%cooperativity
dynamic(1)= g/(1+g)*1/(1+(x2/s)^n)*1/(1+(x3/s)^n)*1/(1+(x4/s)^n);
dynamic(2)= g/(1+g)*1/(1+(x1/w)^n)*1/(1+(x3/s)^n)*1/(1+(x4/s)^n);
dynamic(3)= g/(1+g)*1/(1+(x1/s)^n)*1/(1+(x2/w)^n)*1/(1+(x4/s)^n);
dynamic(4)= g/(1+g)*1/(1+(x1/s)^n)*1/(1+(x2/s)^n)*1/(1+(x3/w)^n);

% static module
n=3;%cooperativity
static(1)= 1/(1+g)*1/(1+(x2/s)^n)*1/(1+(x3/s)^n)*1/(1+(x4/s)^n);
static(2)= 1/(1+g)*1/(1+(x1/s)^n)*1/(1+(x3/s)^n)*1/(1+(x4/s)^n);
static(3)= 1/(1+g)*1/(1+(x1/s)^n)*1/(1+(x2/s)^n)*1/(1+(x4/s)^n);
static(4)= 1/(1+g)*1/(1+(x1/s)^n)*1/(1+(x2/s)^n)*1/(1+(x3/s)^n);

%total gene regulation: dynamic module contribution + static module
%contribution - decay
c=2;
d=3;
dx(1)= c*static(1)+d*dynamic(1)-lambda*x1;
dx(2)= c*static(2)+d*dynamic(2)-lambda*x2;
dx(3)= c*static(3)+d*dynamic(3)-lambda*x3;
dx(4)= c*static(4)+d*dynamic(4)-lambda*x4;

dx=dx';
```

<u>Matlab/Octave code for Movie S4</u>

```
function MovieS4

clear
clc
close all

%Enter which simulation option you would like to run: A, B, C, or D
%These correspond to simulations in Figure 6: Wt, Kr-, kni-, gt-
%and Movie S4 A, B, C, and D
option= 'A';

if option=='A' % WT
    genotype=[1 1 1 1];
elseif option=='B' % Kr-
    genotype=[1 0 1 1];
elseif option=='C' % kni-
    genotype=[1 1 0 1];
elseif option=='D' % gt-
    genotype=[1 1 1 0];
else
    error('Available options are either A, B, C, or D')
end



%Time Axis
t=0:.01:4;

% AP_axis
AP_axis=0:.001:.7;

%solve model
spacetime=zeros(4,length(t),length(AP_axis));
for pos=1:length(AP_axis)
    %initial conditions for Drosophila genes hb, Kr, kni, gt
    initial_conditions=[2*exp(-8*AP_axis(pos)) 0 0 0 genotype];
    [T, x]= ode45(@odefun,t,initial_conditions);
    spacetime(:,:,pos)=x(:,1:4)';
end

%plot solution
close all
for nt=1:length(t)

    %posterior gradient (speed regulator) set up
    g=0.3*exp(-0.2*t(nt))*ones(size(AP_axis));

    set(gca, 'ColorOrder', [86 128 193; 229 51 50; 79 185 118; 219 152 40;
128 128 128 ]/256, 'NextPlot', 'replacechildren');

    plot(AP_axis,[squeeze(spacetime(:,nt,:))' g'],'LineWidth',5)
    axis([0 max(AP_axis) 0 max(max(max(spacetime)))])
```

```
    axis off
    pause(.00000001)
end

%%%%% Model %%%%%%
function dx = odefun(t,x)

x1=x(1);%hb
x2=x(2);%Kr
x3=x(3);%kni
x4=x(4);%gt
genotype=x(5:end);


w=2.5;%Dissociation constant for weak repressions
s=.4;%Dissociation constant for strong repressions
lambda=1;%mRNA decay rate

%posterior gradient (speed regulator) set up
g=0.3*exp(-0.2*t);

%dynamic module
n=5;%cooperativity
dynamic(1)= g/(1+g)*1/(1+(x2/s)^n)*1/(1+(x3/s)^n)*1/(1+(x4/s)^n);
dynamic(2)= g/(1+g)*1/(1+(x1/w)^n)*1/(1+(x3/s)^n)*1/(1+(x4/s)^n);
dynamic(3)= g/(1+g)*1/(1+(x1/s)^n)*1/(1+(x2/w)^n)*1/(1+(x4/s)^n);
dynamic(4)= g/(1+g)*1/(1+(x1/s)^n)*1/(1+(x2/s)^n)*1/(1+(x3/w)^n);

% static module
n=3;%cooperativity
static(1)= 1/(1+g)*1/(1+(x2/s)^n)*1/(1+(x3/s)^n)*1/(1+(x4/s)^n);
static(2)= 1/(1+g)*1/(1+(x1/s)^n)*1/(1+(x3/s)^n)*1/(1+(x4/s)^n);
static(3)= 1/(1+g)*1/(1+(x1/s)^n)*1/(1+(x2/s)^n)*1/(1+(x4/s)^n);
static(4)= 1/(1+g)*1/(1+(x1/s)^n)*1/(1+(x2/s)^n)*1/(1+(x3/s)^n);


%total gene regulation: dynamic module contribution + static module
%contribution - decay
c=2;
d=3;
dx(1)= c*static(1)+d*dynamic(1)-lambda*x1;
dx(2)= c*static(2)+d*dynamic(2)-lambda*x2;
dx(3)= c*static(3)+d*dynamic(3)-lambda*x3;
dx(4)= c*static(4)+d*dynamic(4)-lambda*x4;

dx=dx.*genotype';

%zero change in parameters
dx(5)=0;
dx(6)=0;
dx(7)=0;
dx(8)=0;
dx=dx';
```
